# Interactions between pili affect the outcome of bacterial competition driven by the type VI secretion system

**DOI:** 10.1101/2023.10.25.564063

**Authors:** Simon B. Otto, Richard Servajean, Alexandre Lemopoulos, Anne-Florence Bitbol, Melanie Blokesch

## Abstract

The bacterial type VI secretion system (T6SS) is a widespread, kin-discriminatory weapon capable of shaping microbial communities. Due to the system’s dependency on contact, cellular interactions can lead to either competition or kin protection. Cell-to-cell contact is often accomplished via surface-exposed type IV pili (T4P). In *Vibrio cholerae,* these T4P facilitate specific interactions when the bacteria colonize natural chitinous surfaces. However, it has remained unclear whether and, if so, how these interactions affect the bacterium’s T6SS-mediated killing. In this study, we demonstrate that pilus-mediated interactions can be harnessed to reduce the population of *V. cholerae* under liquid growth conditions in a T6SS-dependent manner. We also show that the naturally occurring diversity of pili determines the likelihood of cell-to-cell contact and, consequently, the extent of T6SS-mediated competition. To determine the factors that enable or hinder the T6SS’s targeted reduction of competitors carrying pili, we developed a physics-grounded computational model for autoaggregation. Collectively, our research demonstrates that T4P involved in cell-to-cell contact can impose a selective burden when *V. cholerae* encounters non-kin cells that possess an active T6SS. Additionally, our study underscores the significance of T4P diversity in protecting closely related individuals from T6SS attacks through autoaggregation and spatial segregation.

## Introduction

The composition of microbial populations has a significant impact on ecological functions and host health [1,2]. Interbacterial interactions are often antagonistic in nature and target closely related species, ultimately influencing microbial populations by aiding in niche colonization and exclusion [3–5]. In order to achieve this, bacteria possess a myriad of weapons, and defence mechanisms [6]. Interactions between species can be achieved through two primary mechanisms: the release of diffusible compounds and contact-dependent interactions [7,8]. Unlike diffusible compounds, which can disperse into the surrounding environment, contact-dependent mechanisms require direct cell-to-cell interaction to effectively deliver toxins [9]. As a result, contact-dependent mechanisms can precisely target neighbouring competitors without the risk of toxin dilution in the surrounding liquid environment [10].

The Type VI secretion system (T6SS) is a widespread molecular apparatus that relies on direct contact to inject toxic effector proteins into target cells [11]. It is estimated to be found in more than 25% of gram-negative bacteria, encompassing both environmental and pathogenic species [12]. The delivery of T6SS toxins has been shown to influence microbial population compositions [13–15]. Importantly, the spatial arrangement of competing cells also impacts the effectiveness of the T6SS [13,16].

The T6SS doesn’t discriminate when targeting cells, owing to its indiscriminate delivery method and toxins that disrupt widely conserved cellular processes [17]. To prevent self-harm and protect kin, T6SS-positive bacteria produce matching pairs of effector and immunity proteins [18]. Yet, bacteria can possess multiple T6SS gene clusters along with variable auxiliary clusters that frequently encode additional effector and immunity proteins [19]. Incompatibility in just one of these effector-immunity pairs can drive T6SS competition [17,20]. Interestingly, targeted cells can also protect themselves through immunity-independent mechanisms, such as through the production of extracellular polysaccharides and surface-attached capsules, allowing them to survive T6SS attacks [21–23]. Furthermore, bacteria have been shown to withstand T6SS challenge, and respond more potently [23–25].

Bacterial autoaggregation is a process that allows bacteria to bind to themselves, often serving as a necessary step in forming biofilms [26,27]. This phenomenon is frequently linked to pathogenicity as it provides protection against external threats, such as phagocytosis [28] and antimicrobial agents [29,30]. Various molecules/structures, often referred to as autoagglutinins, can facilitate this cell-to-cell binding [31]. Here, we specifically focus on Type IV pili (T4P), which are common surface-exposed appendages with diverse functions, including DNA uptake, motility, adhesion, and aggregation [32,33]. T4P’s ability to sense the environment is crucial for the survival, colonization, and virulence of species carrying these pili [34]. For instance, the toxin co-regulated pilus, exclusive to the pandemic lineage of *Vibrio cholerae*, plays a critical role in host and interbacterial cell adhesion, which is vital for pathogenesis [35]. Similarly, T4P mediate interbacterial interactions in other pathogens like *Pseudomonas aeruginosa* and *Neisseria meningitidis* [36,37]. However, this ability to sense the environment and interact with other bacteria might inadvertently lead to unwanted cell-to-cell contact, potentially inviting competition from the T6SS. It is therefore worth noting that many autoagglutinins, including T4P, can modulate their level of aggregation [31,38].

To explore the impact of the T6SS on bacterial behaviour in the context of T4P-mediated cell-to-cell contact, we selected *V. cholerae* as our model organism. This bacterium is notable for possessing both a T6SS and a variable DNA-uptake T4P. Indeed, in *V. cholerae* strains, a single T6SS machinery is responsible for delivering distinct antibacterial effectors. These effectors are encoded within three genetic clusters, including the primary large gene cluster that houses most of the structural components, as well as auxiliary clusters 1 and 2 [19]. In some *V. cholerae* strains, including the current pandemic lineage, there are additional auxiliary clusters that contain extra effector-immunity pairs, although their presence varies [20,39–41]. It is worth noting that in the aquatic environment, the presence of chitin degradation products has a dual effect on *V. cholerae*: it activates both the T6SS machinery and the DNA-uptake T4P as part of the bacterium’s natural competence program. Consequently, T6SS-mediated neighbour predation leads to DNA acquisition, ultimately driving horizontal gene transfer through the process of natural transformation [42–44]. In the current pandemic strains of *V. cholerae*, this activation is orchestrated by the TfoX master regulator once the bacteria reach a high cell density state [42,43,45,46]. In environmental *V. cholerae* isolates, the T6SS machinery is in a state of constant activity [20,47–50], representing an immediate risk for T6SS-associated harm in case cell-to-cell contact in established. Therefore, bacteria must distinguish between nearby individuals before intentionally initiating cell-to-cell contact.

Prior research has shown that the DNA-uptake T4P, present in all *V. cholerae* strains, often self-interacts and that it can distinguish between strains based on the variability of the major pilin protein, PilA [51]. Interestingly, such variability in the major pilin protein has been observed in various species carrying T4P [52–54]. Our hypothesis therefore centres on the potential of T4P to be harnessed for targeted T6SS-mediated bacterial elimination by facilitating specific cell-to-cell contact. Typically, studies exploring T6SS-mediated killing are conducted on agar surfaces at high cell densities, where physical contact is forced due to crowding. In contrast, T4P could enable cell-to-cell contact with particular target cells under non-crowded (e.g., liquid) growth conditions, an idea that we tested in this study. We also sought to investigate the relationship between the natural diversity of PilA and the T6SS, aiming to uncover the strategies bacteria might employ to regulate the risk associated to establishing cell-to-cell contact. Finally, we used simulations to determine the factors that might either enable or impede the predation of T4P-carrying bacteria. Our simulations also emphasize the critical role of spatial organization and rapid lysis in the success of T6SS-mediated targeted depletion. Collectively, our study demonstrates that the natural diversity of T4P plays a pivotal role in regulating the extent of non-kin cell-to-cell contact, thus shaping the ensuing competition driven by the T6SS.

## Results and discussion

### T4P facilitate T6SS-mediated killing by fostering cell-to-cell contact

We hypothesized that T4P-mediated autoaggregation might enhance the essential cell-to-cell contact necessary for T6SS-mediated elimination under liquid conditions. To test our hypothesis, we induced the activity of both T6SS and T4P by artificial induction of the TfoX master regulator [43]. We then cocultured T6SS-competent strains (acting as predator) with T6SS-sensitive target strains (prey). We made the prey strains T6SS-sensitive by deleting the genes encoding the four T6SS effector/immunity protein pairs (Δ4E/I), a modification applied to the pandemic *V. cholerae* strain A1552, which served as the chassis for our research throughout this study. Predator strains were engineered to be either T6SS-competent or rendered non-functional by deleting *vasK*, which encodes a critical component of the T6SS membrane complex [55].

To investigate the role of T4P, we manipulated the major pilin PilA of A1552, a pivotal T4P component. We either kept the encoding gene in its original genetic location (*pilA*(A1552)) or deleted it (Δ*pilA*) in both the predator and prey strains. In all strains, we disabled T4P retraction (Δ*pilT*), which promoted an intensified autoaggregation effect [51]. The phenomenon of amplified autoaggregation due to *pilT* deletion has also been observed in other bacteria, such as *Neisseria meningitidis* [56]. In this study, we employed *pilT* deletion as a practical approach to increase the likelihood of self-interactions between pili. However, it’s worth noting that these enhanced self-interactions may resemble the interactions of T4P on chitin surfaces, where dense networks of pili-pili interactions are well-documented [51]. Unfortunately, performing experiments on chitin surfaces proved to be technically unfeasible for testing the hypotheses outlined above.

Predator and prey strains were distinguishable by the presence of antibiotic/fluorescent markers, which themselves displayed no discernible fitness advantage or disadvantage when compared to a reference strain (Fig. S1A). We then assessed the fitness of the predator strain over the prey strain by conducting bacterial enumerations, followed by the calculation of the selection rate constant using the method introduced by Travisano and Lenski [57]. Positive values of this constant indicate a fitness advantage of the predator strain.

Our results clearly demonstrated a significant fitness advantage for the predator strains when the T4P were functionally active (Fig. 1A). Conversely, when the predator strain had a non-functional T6SS, any fitness advantage was entirely abolished. This suggests that the fitness advantage observed in predator strains is primarily achieved through T6SS-mediated depletion of prey cells.

**Fig. 1.**
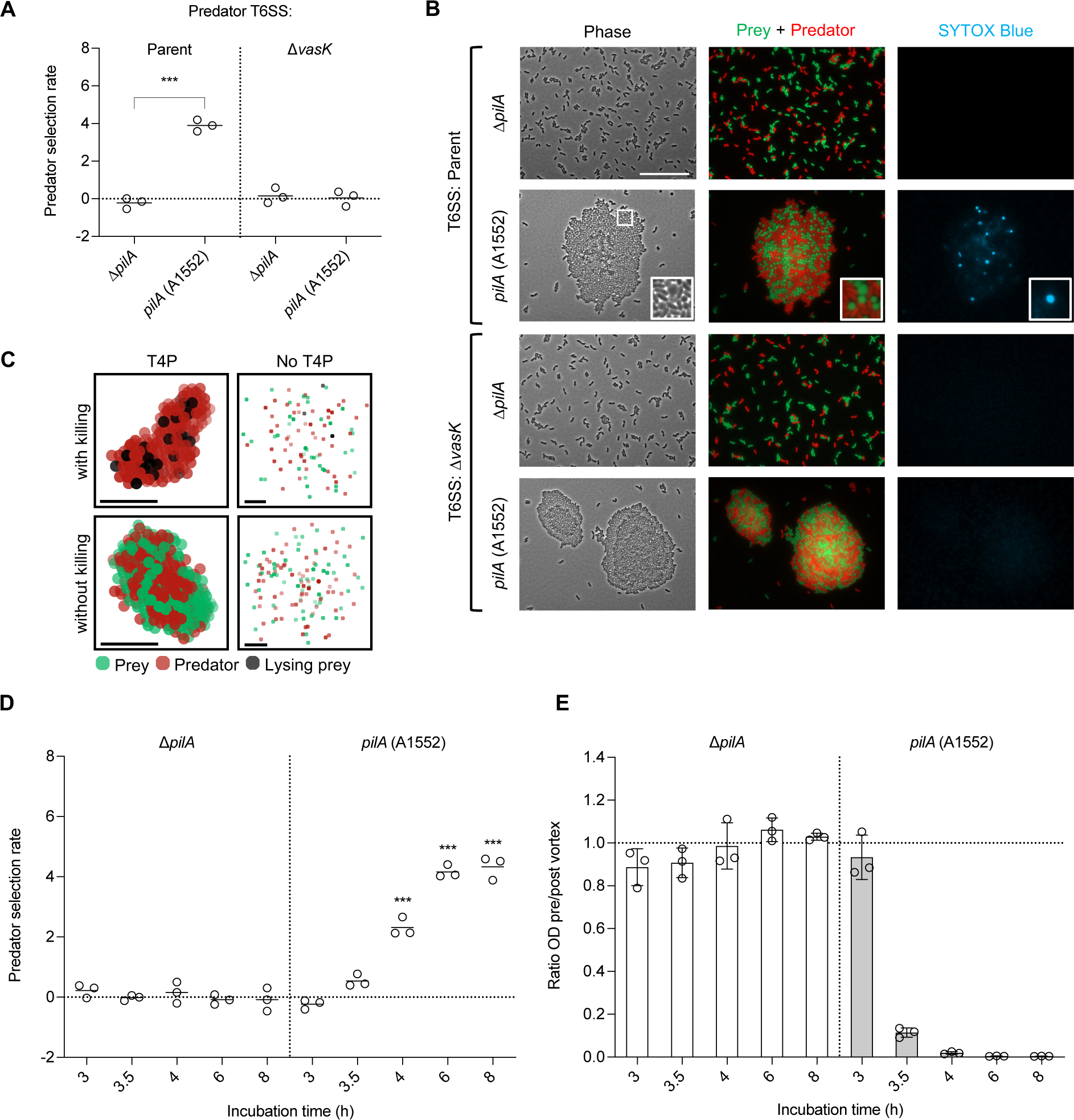
T4P facilitate T6SS-mediated killing in liquid by promoting autoaggregation. **A)** Selection rates of T6SS-competent (Parent) and non-functional (Δ*vasK*) predator strains when co-cultured with T6SS-sensitive (Δ4E/I) prey strains. Both prey and predator strains carry their native major pilin gene (*pilA* (A1552)) or lack it (Δ*pilA*). Selection rates were determined after 6 h of growth. **B)** Representative microscopy images of 4-hour-old co-cultures, corresponding to the experiment in (A). The images include phase contrast (left), a merged view of prey (sfGFP, in green) and predator strains (mCherry, in red) (middle), and SYTOX Blue dead cell stain (in blue) (right). An inset provides a zoomed-in view of an aggregate indicated on the phase channel by a white box. White scalebar: 25 µm. **C)** Zoomed snapshots taken after 1 h from simulations conducted as described in the methods section. The simulations consider T4P self-interactions (left vs. right column) and T6SS-mediated killing (top vs. bottom row) either enabled or disabled. Prey and predator cells are represented by green and red markers, respectively, while lysing cells are indicated by black markers. The black scale bar indicates the length of 5 marker diameters. In the “no T4P” panels, we display the content of an arbitrary cube, of edge about 23 cell diameters, extracted from the system. **D)** Time-course analysis of selection rates of predator strains in co-culture experiments, comparing *pilA*-carrying co-cultures to corresponding *pilA*-deleted co-cultures. **E)** Time-course of co-culture experiments demonstrating the aggregation level at indicated incubation times. The aggregation level is determined by the ratio of the co-culture’s optical density at 600 nm (OD_600_) pre/post-vortex. The horizontal dotted line represents the value around which no aggregation occurs. Graphs show the mean, with error bars giving the standard deviation; circles represent data from three independent experiments. In dot plots, significant differences were determined using one-way ANOVA with Tukey’s post hoc test (\*\*\**P* < 0.001). *P* values are provided in the source data file.

To validate the T6SS-mediated depletion of prey cells, we conducted imaging of the co-cultures, examining the spatial arrangement of prey and predator cells (Fig. 1B). In order to image aggregates, cells were immobilised on an agarose pad and covered with a coverslip (see method section for details). In our observations, we used a cell-impermeable DNA dye (SYTOX Blue) to visualise cells with compromised membrane integrity [58]. What we observed was that the autoaggregation promoted by the T4P effectively facilitated the necessary cell-to-cell contact for T6SS-mediated cell elimination in liquid conditions. In contrast, the absence of functional T4P eliminated the cell-to-cell contact, thereby preventing the killing of T6SS-sensitive strains. Secondly, we noted cell rounding and lysis of T6SS-sensitive prey, a phenomenon predominantly occurring in cells in direct contact with the predator strains (Fig. 1B, inset). This particular trait was dependent on the presence of a T6SS-competent predator strain, with no discernible changes in prey cell morphology when a non-functional T6SS predator strain was utilized. Additionally, the phase contrast images revealed larger aggregates, comprising multiple layers of cells, when a non-functional T6SS predator strain was employed. Thus, these images offer compelling visual evidence of T6SS-mediated prey cell elimination facilitated by the T4P.

To analyse the impact of the different interactions and parameters involved in the T4P-facilitated T6SS-mediated depletion of prey strains, we developed an agent-based model grounded in physical principles, incorporating key biological elements. In this model, prey and predator bacteria were initially randomly distributed at a 1:1 ratio in a three-dimensional 40 × 40 × 40 body-centred cubic lattice modelling the liquid suspension. Subsequently, all bacteria were allowed to move diffusively within the medium, with the ability to divide and to interact through T4P. Predators could execute T6SS-mediated killing of neighbouring prey cells, which then enter a lysing state (for further details, refer to the methods section). By enabling or disabling interactions through T4P and T6SS killing, we were able to qualitatively reproduce the microscopic images observed (Fig. 1C). Just as in the experiments, when T4P were present, mixed aggregates formed, leading to the lysis of prey cells when T6SS killing was enabled. Conversely, consistent with the experimental results, in the absence of T6SS killing, larger aggregates were formed. Moreover, the absence of T4P prevented aggregation, with T6SS killing becoming a rare event. Therefore, our minimal model serves to confirm that prey depletion is significantly enhanced when predator and prey cells adhere via T4P. This heightened contact between neighbouring prey-predator pairs allows for more frequent T6SS killing. Additionally, in the presence of T4P without T6SS killing, bacteria formed mixed aggregates, with a minor tendency to develop homogeneous patches due to cell division. This outcome aligns with the experimental observations, where a low level of patchiness was consistently noted.

In light of these observations, we were keen to delve into the dynamics of contact establishment and the subsequent elimination of prey. To investigate this, we conducted a time-course co-culture experiment in which we determined the selection rate of the predator strain at different incubation times (Fig. 1D). Furthermore, we assessed the levels of aggregation in these co-cultures by comparing the ratio of cells in the solution to those in the settled aggregates (Fig. 1E). What we found was that the fitness advantage of predator strains became evident when compared to a Δ*pilA* co-culture control after 3.5-4 hours of incubation. This synchronization with a preceding rapid aggregation event suggests that the killing of T6SS-sensitive cells occurs shortly after cell-to-cell contact is established. Similar to the experimental observations, simulations exhibit rapid aggregation, followed by T6SS killing (Fig. S2, Movie S1-4). Collectively, our data demonstrate that T4P-mediated aggregation generates enough cell-to-cell contact to facilitate effective T6SS killing, even under conditions where cells are otherwise well-mixed.

Related to this role of T4P in facilitating T6SS-mediated killing in liquid environments, studies have demonstrated that engineered receptor-ligand interactions can lead to the targeted depletion of prey cells within bacterial communities [59]. Similarly, in *Vibrio fischeri*, a putative lipoprotein has been implicated in mediating targeted cell-to-cell contact for T6SS competition in high-viscosity liquid media [10]. It’s worth noting that this lipoprotein’s distribution is limited to bacterial species associated with a marine host and has no homolog in *V. cholerae*. The specificity of target recognition in this case is achieved through an unknown ligand. In various scenarios, not limited to liquid conditions, such as within microcolonies of *Neisseria cinerea*, the expression of T4P by both predator and prey strains heightened the prey’s vulnerability to T6SS attacks in contrast to a non-piliated control. This effect was achieved by preventing the segregation of prey from a T6SS-armed attacker [60]. These findings underscore the potential risk involved in utilizing T4P, as they can serve as potential enhancer of T6SS-mediated competition.

### Naturally occurring diversity of T6SS E/I pairs and T4P pilin alleles

Having demonstrated the potential of T4P to facilitate T6SS-mediated killing in liquid environments, our next objective was to investigate the interplay between the naturally occurring diversity of T6SS and T4P in *V. cholerae*. We were interested in understanding whether there was a selection pressure leading to coevolution between the *pilA* alleles and T6SS compatibility, or if other selective forces influenced the diversity of these two systems. To address this question, we initially constructed a cladogram involving 39 *V. cholerae* genomes, comprising both environmental and patient isolates. We used *Vibrio mimicus* as an outgroup for this analysis (Fig. 2A). To evaluate the diversity of the T6SS, we aligned the nucleotide sequences of the six known T6SS clusters. We then extracted the amino acid sequences of the core immunity proteins. The resulting heatmaps, displaying the percentage identity of T6SS (Figs. S3-S8), were used to group T6SS effector/immunity (E/I) modules into families (sharing over 30% identity) and subfamilies (identical sequences) consistent with methods previously developed [17,61]. This typing approach enabled us to identify six novel T6SS effector/immunity families within the large gene cluster. Additionally, it expanded our understanding of strains carrying the recently discovered auxiliary clusters 4 and 5 [20,40,41] (Fig. 2A).

**Fig. 2.**
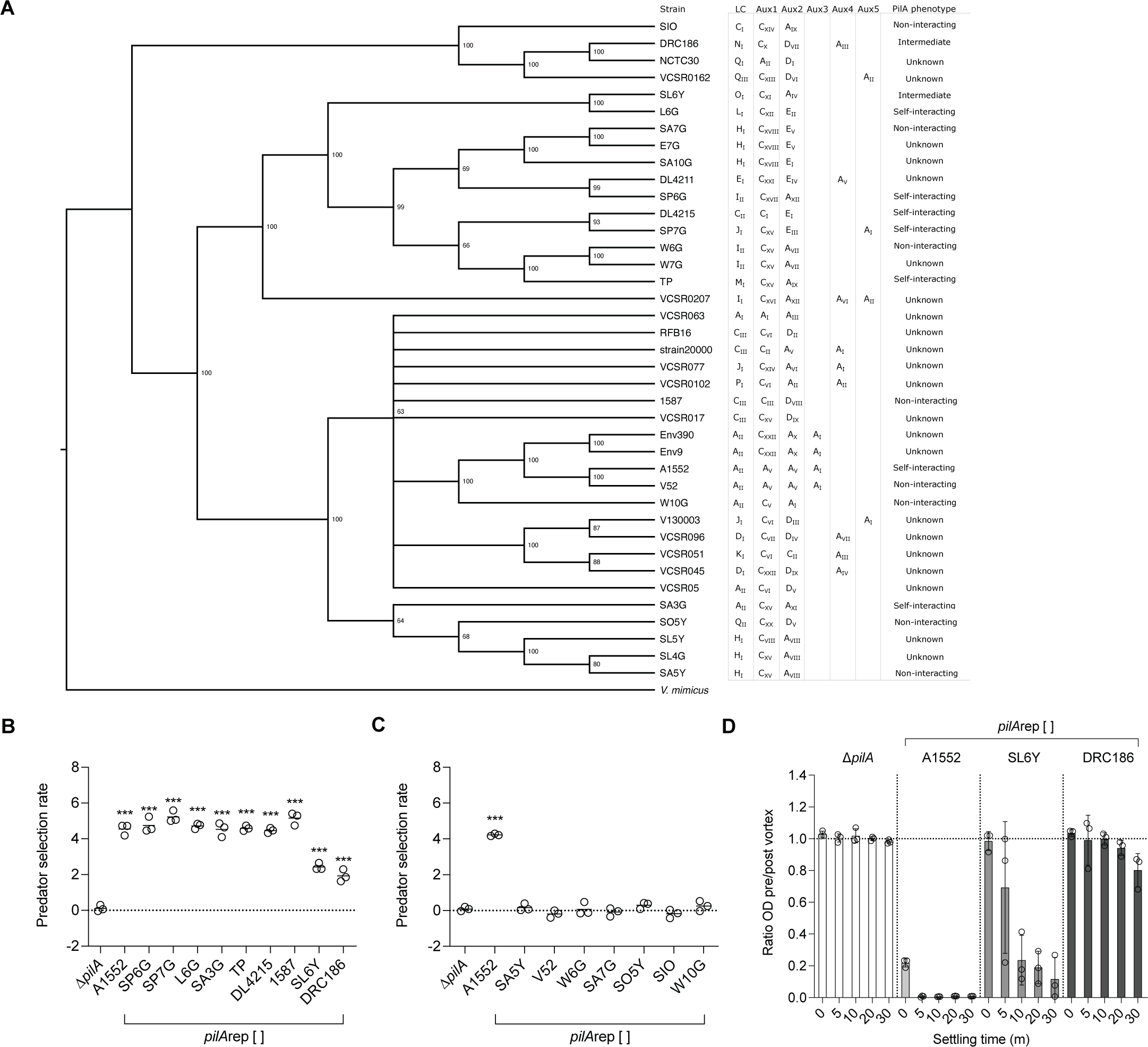
Conservation of T4P-Mediated T6SS Killing Across Self-Interacting PilA Variants. **A)** Cladogram presenting 39 *V. cholerae* strains analysed based on 1186 core genes. *V. mimicus* (ATCC33655) is used as an outgroup. Statistical support is evaluated through 100 bootstraps, with nodes below 60 being collapsed. T6SS effector module families in the large- and auxiliary clusters (LC, Aux 1-5) are specified behind the strain names, followed by the self-interaction ability of T4P. **B)** Bacterial competition assay introducing self-interacting PilA variants into T6SS-competent (predator) and T6SS-sensitive (prey) strains. **C)** Selection rate of non-interacting PilA variants, with self-interacting *pilA*rep[A1552] serving as a positive control. **D)** Aggregation assay assessing the aggregation level of PilA variants SL6Y and DRC186. Cultures are allowed to settle for the specified duration, after which the aggregation level was determined by calculating the ratio of the culture’s OD_600_ before and after vortexing. The horizontal dotted line denotes the ratio around which no aggregation occurs. Δ*pilA* and *pilA*rep[A1552] are designated as negative and positive controls, respectively. Circles in graphs represent independent replicates, while bars/lines indicate the mean, with error bars illustrating the standard deviation. Selection rates of predators were compared to Δ*pilA* for statistical analysis (ANOVA with Tukey’s post hoc test; *** *P* < 0.001). *P* values are provided in the source data file.

The categorization of strains into T6SS families, as depicted to the right of the cladogram in Figure 2A, allowed us to visualize whether they are capable of coexisting, sharing identical T6SS modules, or if they can engage in T6SS-dependent competition. Variety in T6SS families within *V. cholerae* isolates was large, with few T6SS compatible strains. Indeed, previous experimental data among a subset of strains showed limited compatibility between non-kin T6SS systems [20]. Next, we examined the experimentally confirmed self-interaction ability of PilA, which is presented alongside the T6SS modules. We also extracted nucleotide and protein sequences based on genome annotations to construct a heatmap and a PilA cladogram (Fig. S9-10). Notably, the studied PilA variants exhibit high variability and cluster into numerous phylogenetic groups.

Our analysis did not reveal distinct clades for either the T6SS E/I modules or PilA variants (Fig. 2A, S10). While closely related strains occasionally shared similar T6SS E/I modules and PilA variants (for example, strains Env9 and Env390), more often, we observed diversity in both the T6SS E/I modules and PilA variants (such as SP7G and DL4215). Indeed, we did not identify clear-cut markers of coevolution between PilA variants and T6SS compatibility. Furthermore, we observed a lack of congruence between genome-based and *pilA* topologies (Fig. S10) and an average approximately 9% lower GC content for both the T6SS E/I modules and *pilA.* These observations suggest that both *pilA* and T6SS E/I modules can be horizontally acquired, as previously suggested for the latter [17,62]. Horizontal exchange of *pilA* alleles or T6SS E/I modules could thereby alter bacterial competition, as outlined below.

### T4P-facilitated neighbour predation is conserved among self-interacting PilA variants

Subsequently, we assessed whether the capacity of T4P to facilitate T6SS-mediated killing is conserved among different PilA variants. To investigate PilA variability, we introduced *pilA* alleles into the native *pilA* locus of strain A1552, a method previously demonstrated to be fully functional [51]. The predator and prey strains with replaced *pilA* (*pilA*rep) were co-cultured, and their predator selection rates were evaluated (Fig. 2B,C). Our results revealed a positive predator selection rate for PilA variants capable of self-interaction, as compared to a negative control lacking *pilA* (Δ*pilA*; Fig. 2B). Conversely, PilA variants that could not self-interact according to previous work [51] were unable to facilitate T6SS-mediated killing (Fig. 2C).

Interestingly, two PilA variants, from strains SL6Y and DRC186, displayed an intermediate level of prey strain depletion. Both of these PilA variants could support the killing of T6SS-sensitive prey, but the culture tubes appeared more turbid than those with other self-interacting PilA variants that mostly settled to the bottom of the tube. We hypothesized that a weaker aggregation phenotype might allow a subpopulation of targeted prey cells to escape. To explore this, we determined the aggregation levels of these PilA variants and compared them to the strongly self-interacting A1552 PilA variant and the negative Δ*pilA* control (Fig. 2D). While measuring aggregation ratios [51], various settling times were considered to allow for self-interaction of potentially weaker PilA variants. Indeed, both the SL6Y and DRC186 PilA variants exhibited weaker levels of aggregation than the strongly self-interacting A1552 PilA variant. Of these two, the SL6Y variant displayed stronger aggregation during extended settling periods, which are, however, not included in the evaluation of the predator selection rate during co-culture. The major pilin of T4P often exhibits variation [52–54]. This variability can influence the strength of attractive forces between bacterial cells, which can explain our results. Such an effect was seen in *Neisseria gonorrhoeae* [63]. Similarly, the diversity in PilA variants in *Acinetobacter baumannii* was shown to impact the level of T4P self-interaction and its functional specialization [54].

### Pilus-specific T6SS competition by spatial segregation

A significant diversity of PilA variants is naturally present within the non-pandemic *V. cholerae* isolates. We speculated that PilA diversity might provide protection for cells against T6SS competition by enabling specificity in contact establishment. To investigate this, we conducted a co-culture experiment in which we co-cultivated a prey strain with various PilA variants alongside a predator strain containing the pandemic A1552 PilA variant (Fig. 3A). As expected, the presence of PilA diversity negated the fitness advantage of the predator strain, in contrast to the co-culture with a control T4P matching the A1552 PilA variant.

**Fig. 3.**
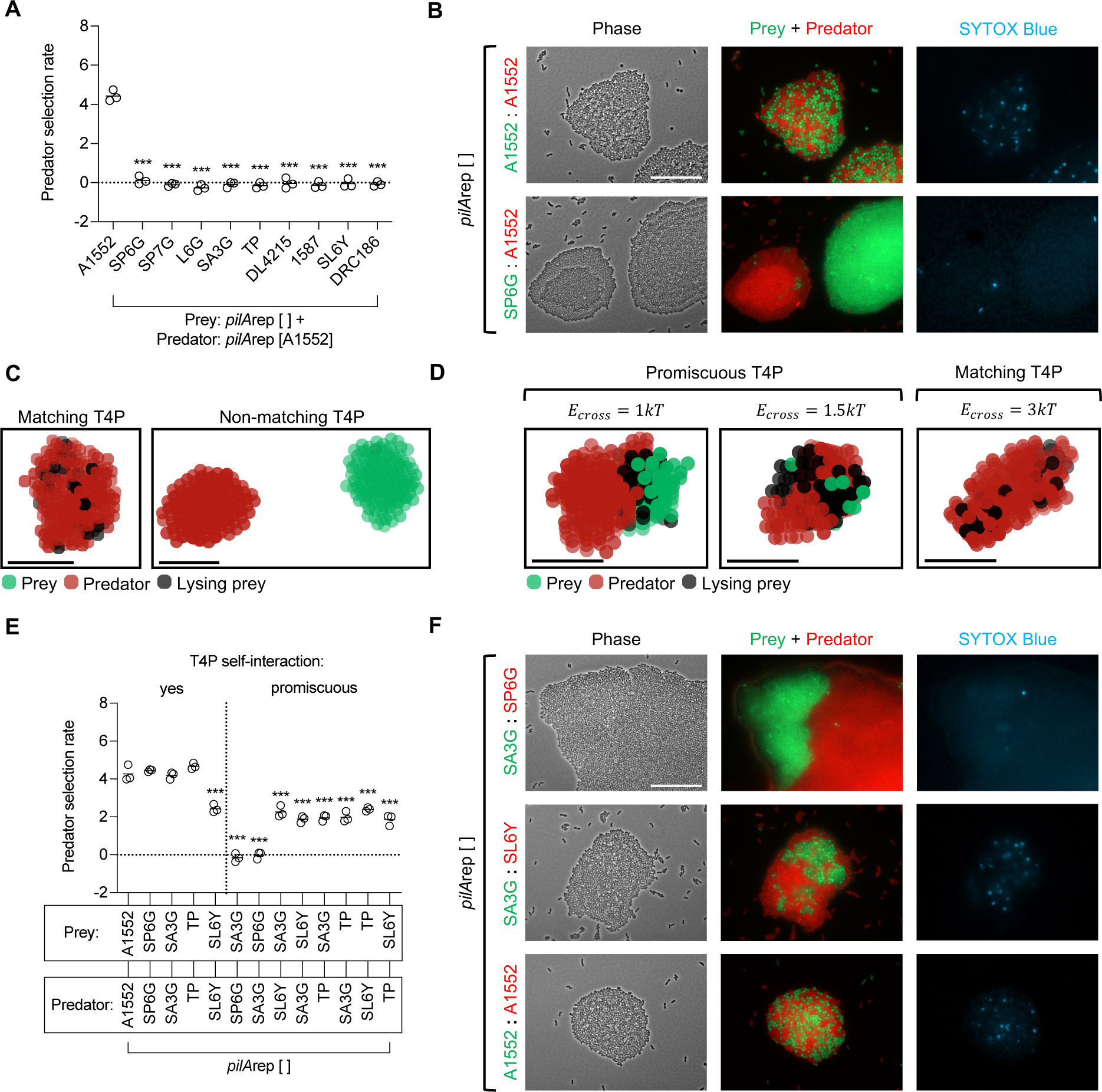
Impact of Pilus Specificity on T6SS Competition and Spatial Segregation. **A)** Selection rate of T6SS-competent (predator) strains competing with T6SS-sensitive (prey) strains, with diverse PilA variants among the prey strains, against a predator strain carrying the A1552 PilA variant. **B)** Representative microscopy images of prey/predator co-cultures, exhibiting either matching (A1552 : A1552) or diverse (SP6G : A1552) PilA variants. Images include phase contrast, a merge of prey (sfGFP, in green) and predator strains (mCherry, in red), and SYTOX Blue dead cell stain (blue). The white scale bar denotes 25 µm. **C)** Snapshots from simulations after 1 h, with varied T4P-mediated binding energy (*E_cross_*) between predators and prey. *E_cross_* was set at 3*kT* for matching T4P and 0 for diverse T4P. **D)** Snapshots from simulations after 1 h, representing various levels of T4P promiscuity obtained by modulating the T4P-mediated binding energy *E_cross_*, compared to a matching T4P control. **E)** Selection rates in prey/predator co-culture experiments, where the carried PilA variants display differing levels of promiscuous T4P self-interactions. **F)** Imaging of co-culture experiments between T6SS-competent (predator) and T6SS-sensitive (prey) strains, featuring PilA variants with varying levels of promiscuousness. The left panel shows phase contrast, the middle panel displays a merge of prey (sfGFP, in green) and predator (mCherry, in red) strains, while the right panel exhibits SYTOX Blue dead cell stain (blue). The white scale bar indicates 25 µm. Circles in graphs represent independent experiments and means are indicated by lines. Statistical tests were compared with the (non-promiscuous) matching *pilA*rep[A1552] control condition using ANOVA with Tukey’s post hoc test (*** *P* < 0.001). For simulations, prey and predators are represented by green and red markers, respectively. Lysing cells are denoted by black markers. The black scale bar indicates the length of 5 marker diameters.

After this discovery, we proceeded to visualize the spatial arrangement of the cells, both with and without PilA diversity (Figs. 3B, S11). While identical T4P led to a well-blended community with numerous interfaces of cell-to-cell contact between the two cell types, T4P diversity facilitated the predominant spatial separation of the two cell types. Consequently, due to the T4P specificity between the pandemic and other self-interacting PilA variants, T4P-mediated T6SS competition with strains carrying diverse *pilA* alleles was circumvented. We therefore conducted simulations that accounted for the presence of non-matching T4P incapable of cross-interaction, in contrast to matching T4P (Fig. 3C, Movie S4-5). Similar to the experimental observations, prey and predator cells with diverse T4P formed distinct aggregates. This prevented encounters between the prey and predator, along with subsequent T6SS-mediated killings, within the timescale considered.

Most PilA variants of *V. cholerae* appear to exhibit highly specific interactions [51]. Interestingly, naturally occurring PilA variants with cross-interaction have also been identified. These promiscuous PilA variants could facilitate cell-to-cell contact between potential T6SS competitors and invite T6SS competition. We hypothesized that the potential fitness disadvantage of T6SS competition could be offset by preferential binding to kin. To further investigate this, we simulated prey and predator interactions, in which we reduced the cross-interaction T4P binding energy (*E_cross_*) between prey and predator, making cross-interaction weaker than self-interaction (Fig. 3D, Movie S6). These simulations resulted in patchy aggregates. Increasing *E_cross_* within this range led to increased mixing within aggregates, providing more boundary interfaces between prey and predator, until the cross-interaction T4P binding energy equalled the self-interaction T4P binding energy between two prey cells or two predators, resulting in an aggregate simulating matching T4P. Simulating preferential binding to kin restricted T6SS killing predominantly to the boundary interfaces between patches, thanks to the partial segregation effect arising from weaker interactions across T4P variants.

Next, we performed coculture experiments to evaluate the predator selection rate of all known promiscuous interactions, as well as their conforming counterparts (Fig. 3E). Simultaneously, we determined the spatial organization of the cells for all promiscuous interactions (Figs. 3F, S12). Compared to the matching A1552 PilA variant condition, we observed two distinct phenotypes. The promiscuity between SA3G and SP6G PilA variants resulted in low-level mixing, with sparse non-kin cell-to-cell contact at boundary interfaces. This led to no fitness advantages for T6SS competent cells at the population level, as evident from the determined predator selection rate. For the other promiscuous PilA variant combinations, we observed an intermediate phenotype. Segregated groups of cell types were observed, leading to a subpopulation of cells engaging in non-kin cell-to-cell contact, while others established cell-to-cell contact between kin cells. This matches the predictions from our model, since the simulations that modelled preferential binding to kin produced aggregates of similar morphology to both types of promiscuous T4P, depending on the exact value of the cross-interaction T4P binding energy (*E_cross_*). This suggests that weaker interactions between promiscuous T4P variants are responsible for the observed spatial organization patterns. The resulting patchiness leads to a reduction of T6SS competition interfaces compared to the case with matching pili, and subdivides the potential fitness advantages or disadvantages of T6SS competition to a subset of the population, resembling a bet-hedging strategy. Alternatively, intrinsically modulating the strength or ability of T4P to self-interact (e.g. SL6Y/SA5Y) would reduce the number of all T4P-facilitated cell-to-cell contacts but may affect the functional specialization of the strain. Lastly, the predominant PilA variant specificity found in naturally occurring isolates suggests a selective burden for ambiguous T4P. Although we demonstrate that T6SS competition could apply a selective burden for pilus conformity, we suggest that diversity in both T6SS and PilA is also driven by factors independent of their interplay. Namely, T4P-independent T6SS competition, potentially caused through proximity by crowding, could also drive the diversity observed in T6SS E/I modules, while the PilA variability might reflect the selection pressure by other stressors, such as phage predation.

### Spatial assortment and lysis time dictate T6SS-mediated target cell depletion

With evidence of T4P’s potential in facilitating T6SS depletion, we aimed to explore the key factors influencing its effectiveness using our agent-based model. Considering the observed ability of T4P to regulate non-kin cell-to-cell contact, we assessed the impact of cross-interaction T4P binding energy on mixing levels (Fig. 4A). To quantitatively analyse the mixing within aggregates, we evaluated assortment, which compares the number of observed adjacent prey-predator pairs to its maximum expected value in the case of random mixing (see methods). Similar to the experimental observations, our simulations demonstrate that a non-zero cross-interaction T4P binding energy (*E_cross_*) between prey and predator increases assortment, thereby promoting mixing. Higher cross-interaction T4P binding energy between prey and predator results in the formation of more mixed aggregates, with larger *E_cross_* values leading to increased spatial assortment. Moreover, we noted a stronger long-term gradual rise in assortment over time in the presence of T6SS killing compared to scenarios without T6SS killing. The action of T6SS results in the lysis of prey cells that do not undergo division, thus reducing the division-induced tendency of aggregates to develop distinct patches. Under infinite time, all T6SS killing simulations would eventually become uniform, consisting solely of predators, corresponding to fixation of the predator type. However, it is crucial to highlight that our simulations do not account for the potential escape of divided prey cells into new niches. In natural settings, the observed delay in spatial assortment for lower values of *E_cross_* could potentially be exploited by the T6SS-sensitive prey.

**Fig. 4.**
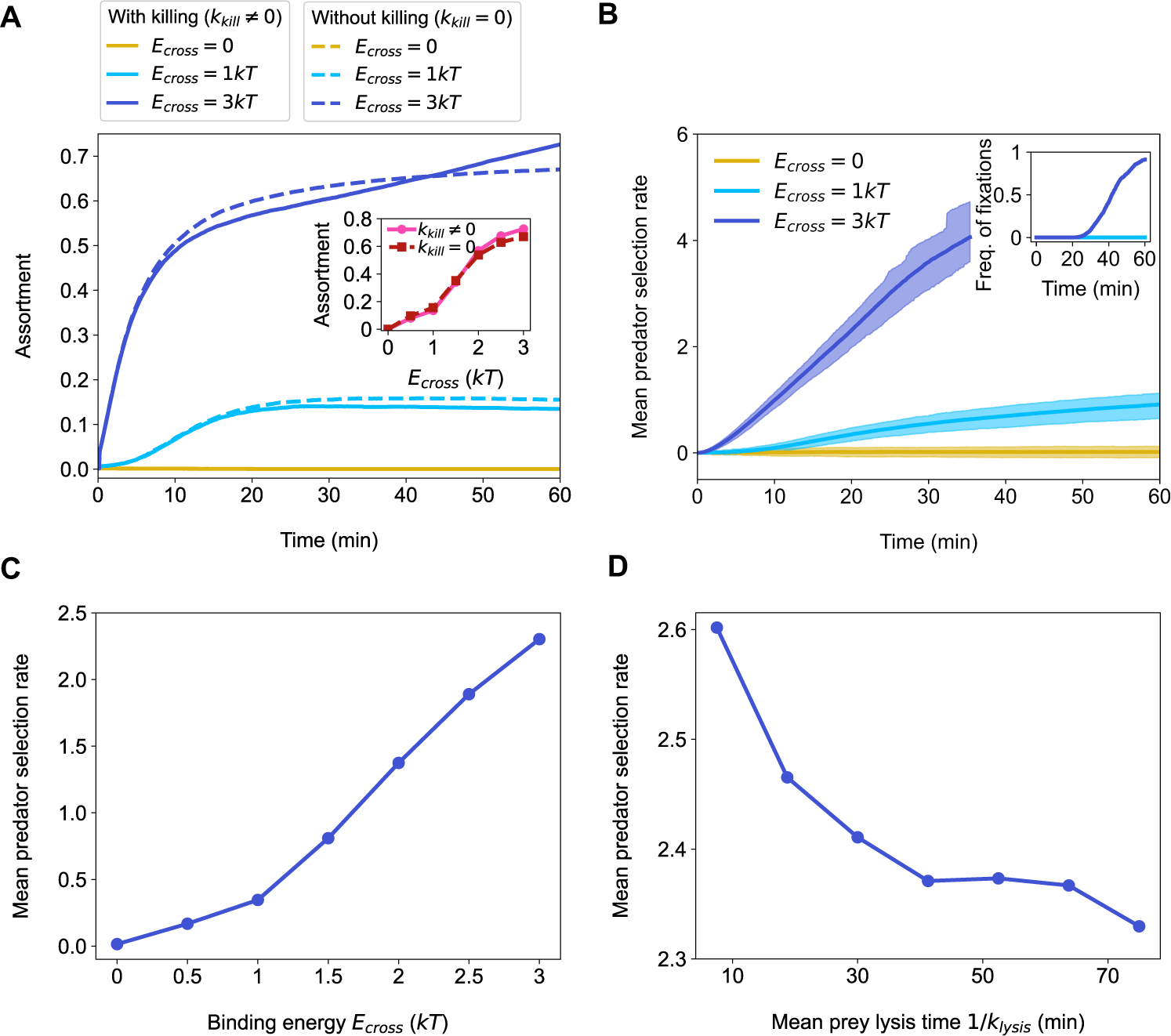
Influence of Spatial Assortment and Lysis Time on T6SS-Mediated Target Depletion. **A)** The impact of the attractive interaction between prey and predators on mixing over time. The main panel displays the assortment between prey and predators over time for different values of the T4P binding energy *E_cross_* between a prey and a predator. Solid lines represent simulations with killing (*k_kill_* = 6.25*h*^-1^), while dashed lines depict scenarios without killing (*k_kill_* = 0). Assortment is determined by comparing the number of adjacent prey-predator pairs to its maximum expected value, considering lysing cells as prey (see methods). The inset graph shows assortment after 1 h versus *E_cross_*. **B)** Evolution of selection rate over time. The main panel depicts the mean selection rate versus time for various *E_cross_* values. The inset showcases the frequency of fixation events, where only predators remain, and prey are completely depleted, across the replicate simulations. When fixation occurs in a replicate, its selection rate diverges, and is thus excluded from the calculation of the mean selection rate. To address potential bias, the results are displayed until the frequency of fixations reaches 0.25 for *E_cross_* = 3*kT*. The shaded areas represent the interquartile ranges. **C)** The influence of attraction between prey and predators on the resulting predator selection rate. The graph displays the mean selection rate versus *E_cross_* after 20 minutes of simulation. **D)** The effect of lysis time on prey depletion. The mean predator selection rate is shown versus the mean lysis time (1/*k_lysis_*) after 20 minutes of simulation with *E_cross_* = 3*kT*. When a fixation occurs in a replicate simulation, the corresponding selection rate diverges, and is therefore not considered in the calculation of the mean selection rate. Fixation events were observed in less than 0.5% of replicate simulations. All results are averaged over 10^3^ replicate simulations.

Having quantified how T4P cross-interactions shape spatial assortment, we aimed to determine the impact of these interactions on the predator selection rate (Fig. 4B). We observed a rapid increase in the selection rate with time, greatly influenced by the value of the prey-predator T4P binding energy. When *E_cross_* = 0, this increase was minimal within the time scale considered. Therefore, strong T4P-mediated attraction between prey and predator plays a crucial role in promoting T6SS competition, a condition achieved by matching T4P, and modulated by promiscuous T4P. We observed a positive relationship between the predator selection rate and *E_cross_*, with T6SS competition being enhanced as prey-predator attraction increases towards ambiguity (Fig. 4C). Matching T4P would ensure a strong binding energy between cell types, leading to the subsequent T6SS depletion of target strains, while diverse T4P would hinder T6SS competition, depending on the level of promiscuity of the PilA variant. Through PilA diversity, spatial assortment is modulated, dictating the dynamics of T6SS-mediated depletion of target strains by altering the level of T6SS competition interfaces. Indeed, autoaggregation facilitated by self-interacting identical T4P could function as a defence strategy of *V. cholerae* against T6SS invaders.

Interestingly, the reduction in T6SS competition interfaces observed in the simulations and imaging of promiscuous T4P resulted in lysing cells forming a barrier between cell types. This phenomenon, known as the corpse barrier effect [64], could significantly influence the dynamics of prey depletion. To further investigate this, we determined the predator selection rate with various lysis times (Fig. 4D) [65]. We observed a negative relationship between lysis time and predator selection rate. Consistent with the findings of Smith and colleagues [64], the corpse barrier effect impedes T6SS competition by obstructing predators from reaching new targets. Predator cells equipped with fast-acting lysing effector modules could overcome this barrier, which might be a critical factor in considering target depletion. In *V. cholerae*, strain W10G, carrying two pandemic-like A-type E/I modules in the large and aux2 clusters, has been shown to exhibit strong killing activity against other environmental *V. cholerae* strains [20]. Moreover, *Vibrio coralliilyticus*, while not resistant to *V. cholerae* T6SS attacks, can withstand T6SS challenge and deplete pandemic *V. cholerae* through T6SS killing [25]. Lastly, spatial assortment could prevent the formation of a corpse barrier, as demonstrated in our experiments. Mixing between predator and prey cells, promoted by matching T4P, increases T6SS competition interfaces. However, it remains to be investigated whether this mixing occurs under host or environmental conditions.

Thus, an important question for future research will be to comprehend whether under natural conditions, T4P could serve as a means for the recruitment of predator cells. This exploration could also yield further insights into the consequences of T6SS and T4P on the local enrichment of pathogenic or environmental strains, as well as PilA variant specialization for *V. cholerae*, specifically. Nonetheless, the swift capture and subsequent elimination of pandemic *V. cholerae* through pilus conformity observed in our experiments have demonstrated the potential for the targeted depletion of T4P-carrying species.

The critical role of self-interacting T4P in the colonization of pathogen hosts and environmental niches may contribute to the maintenance of the targeted receptor. For instance, in *V. cholerae*, the conservation of the strongly self-interacting PilA variant within the pandemic lineage suggests its significance in the aquatic environment and/or during human transmission [51]. Additionally, the preservation of T4P specificity might prevent T4P-facilitated competition with T6SS-carrying competitors. In contrast, the self-interacting toxin co-regulated pilus, exclusive to the *V. cholerae* pandemic lineage, does not display target specificity between major pilin variants [51,66]. It is important to note that the T6SS E/I modules of pandemic strains are identical, rendering them T6SS compatible. Finally, considering the natural occurrence of diverse self-interacting T4P in various pathogens [31,53,54,67], the investigation of targeted depletion using T4P should be pursued further.

## Materials and methods

### Bacterial strains and growth conditions

The bacterial strains and plasmids used in this study are listed in Supplementary Table 1. O1 El Tor strain A1552 was used as a genome-sequenced representative of the ongoing seventh cholera pandemic [68,69]. Unless otherwise specified, bacteria were cultured in lysogeny broth (LB) (Carl Roth, Switzerland) or on LB agar plates. The LB medium was supplemented with 1 mM CaCl_2_ and 5 mM MgCl_2_ to counteract LB batch-to-batch variability in aggregation [51]. Indeed, LB medium is often low in divalent cations [70], and CaCl_2_ and MgCl_2_ concentration can be vastly different between batches/different producers [71]. Notably, the CaCl_2_ and MgCl_2_ concentrations present in seawater are, on average, considerably higher than those supplemented in this study [72]. Despite this, the addition of divalent cations has been found to have no effect on T6SS secretion, although it has been observed to impact the conditional efficacy of certain T6SS effectors [71,73].

Cultures were induced with 0.2% L-arabinose to promote the expression of *P*_BAD-_driven genes that were carried by a mini-Tn7 transposon [74] integrated on the chromosome. Post-bacterial mating, *Escherichia coli* cells were counter-selected using Thiosulfate citrate bile salts sucrose (TCBS, Sigma-Aldrich) agar. SacB-based counter-selection was carried out on NaCl-free LB agar supplemented with 10% sucrose. Various antibiotics such as ampicillin (Amp, 100 μg ml^-1^), gentamicin (Gent, 50 μg ml^−1^), kanamycin (Kan, 75 μg ml−1), or spectinomycin (Spec, 200 μg ml^−1^) were added when necessary. Prior to each experiment, overnight cultures were adjusted to an optical density at 600nm (OD_600_) of 2.0 and if required, mixed in a 1:1 ratio, before being back-diluted 1:100 in fresh LB medium. These cultures were incubated in 14 ml round-bottomed polystyrene test tubes (Falcon, Corning) on a carousel style rotary wheel (40 rpm) at 30°C.

### Bacterial strain engineering

Standard methods were used for DNA manipulations and molecular cloning [75]. All genetically engineered strains were verified by PCR, Sanger sequenced by Microsynth AG and analysed using SnapGene (v. 4.3.11). Genetic engineering of *V. cholerae* was done by natural transformation followed by FLP recombination (TransFLP) [76–78], tri-parental mating [79] or allelic exchange using the counter-selectable plasmid pGP704-Sac28 [80].

### Bacterial competition assays

The selection rate of predator strains was evaluated in co-culture experiments using the specified predator-prey strain pairs. Cultures were typically grown for 6 h, followed by a wash step in PBS and subsequent vortexing for 10 min at maximum speed to disperse the aggregates into single cells. Right after dispersion, cells were serially diluted in PBS, and both prey and predator strains were counted on selective antibiotic-containing plates.

The selection rate of the predator was calculated as the difference in the Malthusian parameters of both strains: *r* = ln*[N_predator_* (*t* = 1)/*N_predator_* (*t* = 0)] – ln*[N_prey_*(*t* = 1)/*N_prey_*(*t* = 0)], where *t* = 1 is the numerical density (*N*) at the end of the experiment [57].

The relative fitness of the strains carrying antibiotic/fluorescent markers was determined in co-culture experiments using A1552Δ*lacZ* as the reference strain, following previously established methods [81]. Overnight cultures were prepared as described earlier, but back-diluted to an OD_600_ of approximately 0.0016 in fresh LB medium and grown without antibiotics for 8 h. At the beginning and end of the experiment, the proportions of blue (tested strain) and white (reference strain) colonies were counted through serial dilution in PBS, followed by plating on LB agar plates supplemented with 5-bromo-4-chloro-3-indolyl β-d-galactopyranoside (X-gal; 40 μg ml^-1^).

The relative fitness was calculated as the ratio of the Malthusian parameters of blue over white colonies: *W* = ln*[N_blue_*(*t* = 1)/*N_blue_*(*t* = 0)] / ln*[N_white_*(*t* = 1)/*N_white_*(*t* = 0)], where *t* = 1 is the numerical density (*N*) at the end of the experiment [82].

### Bacterial imaging through light microscopy

To visualise aggregates, overnight cultures were back-diluted, as mentioned previously, and were grown for 4 h. The cells were immobilized by mounting them on slides coated with an agarose pad (1.2% wt/vol in PBS), covered with a coverslip, and imaged using a Zeiss Axio Imager M2 epifluorescence microscope with an AxioCam MRm camera, controlled by Zeiss Zen software (ZEN 2.6 blue edition). Images were captured using a Plan-Apochromat ×100/1.4-NA Ph3 oil objective illuminated by an HXP120 lamp and were analysed and prepared for publication using Fiji [83]. To stain lysed cells, the agarose pad was supplemented with 0.5 µM SYTOX Blue Nucleic Acid Stain (Thermo Fisher Scientific) as previously described [58].

### Bacterial aggregation assay

Aggregation assays were conducted following the established protocol [51]. Bacteria were grown for 6 h, and unless specified allowed to settle for 30 minutes, or 5 minutes for the time-course experiment (Fig. 1E). The level of aggregation was determined by measuring the OD_600_ before and after vortexing (vortexed at maximum speed for approximately 5 sec), during which the settled aggregates were resuspended into the solution.

### Computational model development and testing

An agent-based model was developed on a lattice, grounded in physical principles and incorporating the key biological components of the system. The focus was directed towards the events subsequent to the induction of T4P and T6SS, precisely from the 3-hour mark in the experiments. Stochastic simulations of the model were conducted, employing experimentally measured parameter values whenever available. The model was simulated on a lattice in both two and three dimensions (Fig. S13), with primary emphasis placed on the three-dimensional case in the main text, owing to its closer approximation to real-world conditions. In the three-dimensional case, each cell sits on a site of a body-centred cubic lattice. The following section outlines the fundamental components of the model.

#### Division

In our model, a cell can divide with rate *k_div_*, matching the experimentally measured value, if at least one of its neighbouring sites on the lattice is empty. The offspring is identical to its parent cell and is placed on a randomly chosen empty neighbouring site. In our experiments, the cell division rate was determined to be *k_div_* = 1.58*h^-^*^1^. The growth rate was calculated using the formula *k_div_* = ln[(*OD*_2_/*OD*_1_)/(*T*_2_ -*T*_1_)], where *OD* represents the optical density and *T* represents the incubation time. The values 1 and 2 correspond to the start and end, respectively, of the linear portion of the optical density curve of the wild-type *V. cholerae* grown under the specified growth conditions. Exact measurements can be found in the source data file.

#### Transport

In our experimental setup, the bacteria are placed in test tubes and subjected to agitation through rotation. This is relevant due to the turbulent flow observed in the natural habitat of *V. cholerae*, predominantly in oceans, estuaries, and rivers [84–86]. In such a turbulent regime, passive transport by the medium can be modelled by eddy diffusion. In addition, *V. cholerae* bacteria can actively swim, but the agitation of the medium prevents any substantial gradient that might bias their motion via chemotaxis or quorum sensing. Thus, their active swimming motion may also be simply modelled by diffusion [87]. We therefore model transport through an effective diffusion coefficient *D*, which incorporates both passive and active transport.

Eddy diffusion coefficients are challenging to measure as they are contingent on local flow velocity and the sizes of eddies [86]. However, they are usually significantly larger than the molecular diffusion coefficient *D_mol_* = 5×10^-13^*m*^2^⋅*s*^-1^, obtained via the Stokes-Einstein equation *D_mol_* = *kT* / (6*πηR*), where *k* is the Boltzmann constant, *T* = 300 *K* is the absolute temperature, *η* = 8×10^-4^*Pa*. *s* denotes the dynamic viscosity of water and *R* is the effective radius of bacteria, i.e. the radius of a sphere with the experimentally-measured volume of a wildtype *V. cholerae* bacterium [88]. Active diffusion coefficients associated to swimming can be expressed from the properties of bacterial swimming trajectories [87], and are of the order of 10^-11^*m*^2^⋅*s*^-1^ for *Escherichia coli* run and tumble motion. In our simulations, we adopted a phenomenological value of *D* = 3×10^-12^*m*^2^⋅*s*^-1^, which was found to reproduce the experimentally observed aggregate formation in the absence of killing by T6SS. However, we note that there is uncertainty on this value. For instance, *V. cholerae* was recently found to swim faster than *E. coli* [89], which could yield a larger effective diffusion coefficient. An increased diffusion coefficient should mainly accelerate cluster formation – and would make simulations more computationally demanding. Importantly, previous work had shown that aggregate formation is maintained in non-motile *V. cholerae* [51].

In addition to individual bacterial cells, aggregates of bacteria bound by T4P interactions may also diffuse as a single unit. We include this effect for completeness, but it is worth noting that there are various possible detailed choices for its implementation (Are neighbouring non-bound bacteria pushed by a moving aggregate? Are they pulled by it? Can aggregates break into large blocks? Can they merge? Is their effective diffusion coefficient the same as for single bacteria? For simplicity we answered yes, no, no, yes and yes to these questions). Given these complications, simulations were also conducted without considering any aggregate diffusion. Figure S14 demonstrates that simulations with and without aggregate diffusion exhibit the same phenomenology in the case of diverse T4P, and we also obtained similar results in other cases. Therefore, this effect or its variants do not influence our conclusions.

In practice, in our lattice model, diffusion is implemented via bacteria hopping randomly to any free neighbouring site, with a rate *k_hop_* which is derived from the diffusion coefficient: *k_hop_* = *dD*/(2*zR*^2^), where *d* = 3 is the dimension considered (3D here), *z* = 8 is the number of nearest neighbours per site in the body-centred cubic lattice, and *R* is the effective radius of a bacteria (see above). Since each site has 8 neighbours, the total hopping rate of a bacteria is 8*k_hop_*, if all its neighbouring sites are free.

#### Interactions via T4P

The effect of T4P is modelled as an attractive interaction between neighbouring bacteria on the lattice. Considering that the T4P of prey and predators may differ, three types of interactions are considered, each with potentially different binding energies: *E_prey_* between two prey cells, *E_predator_* between two predators and *E_cross_* between a prey and a predator. Importantly, these interactions have an impact on the ability of individual bacteria to diffuse. Qualitatively, a bacterium bound to many others will be less likely to move away, due to the requirement of detaching from its neighbours. To model this, we assume a dynamics that ensures detailed balance for these moves [90]. The hopping rate of a prey to any free neighbouring site is *nk_hop_exp*(-*n_prey_E_prey_* - *n_predator_E_cross_*) where *k_hop_* is the baseline hopping rate of a freely diffusing bacterium, while *n* is the number of free neighbouring sites, *n_prey_* the number of neighbouring prey and *n_predator_* the number of neighbouring predators. A similar formula can be written for a predator.

We are not aware of precise measurements of the binding energy associated with T4P in *V. cholerae*. However, the involved interactions are reversible protein-protein interactions, and thus, we expect them to be on the order of a few *kT*, where *kT* is the scale of thermal fluctuations (*k* being the Boltzmann constant and *T* the absolute temperature). Because T4P binding energies are not precisely known, we varied them in our simulations [91]. The first key point is that they need to be strong enough to ensure effective aggregation of prey and predator separately, as observed in the experiments. Indeed, as shown in Figure S15, in a system where there is only this attraction and transport (no division, no killing), aggregation occurs above 2*kT* in 2D and 2.5*kT* in 3D at the densities we considered. These thresholds correspond to the liquid-gas phase transition in a lattice fluid, both in 2D [92] and 3D [93], and they are in good agreement with theoretical mean-field calculations. Therefore, we choose *E_prey_* and *E_predator_* above these thresholds, while not exceeding a few *kT*. In practice we take 3*kT*. Then *E_cross_* should be either the same for matching T4P, or smaller otherwise, and we vary it in the latter case.

#### Killing by T6SS

A predator can kill a neighbouring prey at rate *k_kill_*. Once a killing event occurs, the prey enters a lysing state, and is removed from the system at a certain rate *k_lysis_*. For simplicity, it is assumed that, while lysing, a prey moves and interacts with other bacteria in the same way it did before being killed. However, it is unable to divide. The values *k_kill_* = 6.25*h*^-1^ (total firing rate *k_fire_* = 50*h*^-1^ divided by the 8 possible firing directions in the lattice) and *k_lysis_* = 75*min*^-1^ were chosen based on the ranges found in the literature [64,65].

#### Initial density

In experiments, the average initial inoculum comprises 10^7^ bacteria in 2 mL of solution, and the induction of T4P and T6SS production continues for 3 hours until the observation of aggregates at the 3.5-hour time point (Fig. 1E). Therefore, assuming exponential growth at a rate 1.58*h*^-1^, the density at the onset of aggregation is estimated to be around 10^9^ bacteria per mL. In our 3D simulations, a 40×40×40 body-centred cubic lattice was considered, and the initial population consisted of 100 bacteria (comprising half prey and half predators as in the experiments), resulting in approximately 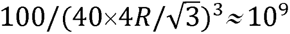 bacteria per mL, where *R* represents the effective radius of a bacterium (see above).

#### Simulation methods

The simulations are conducted using a kinetic Monte Carlo algorithm [90]. Time is discretized with a timestep chosen to ensure that it is unlikely that substantially more than one event occurs within a step, typically on the order of 1 microsecond. Periodic boundary conditions are applied.

#### Assortment: quantitative characterization of mixing within aggregates

Denoting prey by A and predators by B, we define assortment as *n_AB_*/*n_AB, max_*, where *n_AB_* is the number of adjacent prey-predator pairs, while *n_AB_*_,*max*_ is its maximum expected value, obtained if all bacteria were randomly mixed and in the bulk of an aggregate. The latter is *n_AB_*_,*max*_ = *n_A_*×8×*n_B_*/(*n_A_* + *n_B_*), namely the number of prey, times the number of neighbouring sites it has in the lattice (8), times the probability that a neighbouring site is occupied by a predator, assuming that all neighbouring sites are occupied (which is the case in the aggregate bulk), and that prey and predators are randomly mixed. Note that lysing prey are counted as prey in the calculation of assortment.

### Bioinformatical analysis and phylogeny

*Vibrio* spp. genomes used in this study are detailed in Supplementary Table 2. Any genomes that were not previously annotated were annotated using the prokaryotic genome annotation pipeline (PGAP, 2022-10-03. build6384) [94]. To reconstruct the evolutionary history of the studied strains, we first assembled their pangenome along with a *V. mimicus* strain. We used PPanGGOLiN (v. 1.2.74) [95] and provided both sequences and annotation to the program, while the rest of the parameters were set to default. 1186 core genes were identified across the strains. We provided the core genes to Modelfinder [96] to assess gene-specific optimal evolutionary models. Finally, the phylogenetic reconstruction was performed using iqtree2 (v. 2.2.0) [97]. *V. mimicus* was set as the outgroup and 100 bootstraps were computed in iqtree2. Nodes with bootstrap values under 60 were collapsed using the collapseUnsupportedEdges function in the ips package (v. 0.0.11, R environment) [98,99]. GC content was analysed by SnapGene (v. 4.3.11).

T6SS clusters in each genome were identified through blast analysis. A nucleotide database was created using all the studied genomes by employing the makeblastdb command (v. 2.12) with default parameters. The conserved flanking genes for each T6SS cluster were selected based on their known arrangement. These genes were then used as boundaries to identify the T6SS clusters within the genomes. The A1552 sequences were used as query sequences (blastn, default parameters, e-value 1e-10) to detect each respective cluster in the other strains, including the large cluster and auxiliary clusters 1 to 3. For auxiliary clusters 4 and 5, we utilized the same approach using the sequences from *V. cholerae* strain S12 [40] and *V. cholerae* strain BC1071 [41], respectively, as query.

To assess the different families and subfamilies of individual clusters, we selected core immunity proteins for each cluster. These immunity protein sequences were then aligned using muscle (v. 5.1.osx64, default parameters). The hierarchical clustering and the identity matrices for each T6SS cluster were computed in an *R* environment (v. 4.2.1), using the filter.identity function (cutoff = 0.3) in bio3d package (v. 2.4-4). Heatmaps were then visualized with the pheatmap function (v. 1.0.12).

The PilA nucleotide and protein sequences were collected based on the genome annotations for each strain. A phylogenetic tree of the *pilA* nucleotide sequences of the studied strains was reconstructed using iqtree2 (v. 2.2.0), and its statistical relevance was asserted with 100 bootstraps. The best model of evolution was determined using ModelFinder. Nodes with bootstrap values below 60 were collapsed using the collapseUnsupportedEdges function. The protein sequences were aligned using muscle, and the identity matrix was obtained in the *R* environment through the seqidentity function in the bio3d package (v. 2.4-4). The resulting heatmap was visualised with the pheatmap function (v. 1.0.12).

### Statistics and reproducibility

All data represent the outcome of three independent biological experiments, demonstrating consistent results. Bar graphs illustrate the mean value, with error bars denoting the standard deviation. For scatter dot plots, lines represent the mean value. Statistical analyses were conducted using GraphPad Prism (v. 9.1.2). Differences were assessed using one-way ANOVA (α = 0.05) and adjusted for multiple comparisons by the Tukey post hoc test or by a two-tailed Student’s t-test (α = 0.05), when appropriate.

## Data availability

Source data are provided with this paper.

## Code availability

All simulation codes are available on GitHub: https://github.com/Bitbol-Lab/T4P-T6SS-interplay

## Acknowledgements

We thank M. Jaskólska for provision of the codon-optimized mCherry construct, M. Jaskólska & David W. Adams for their contribution to the mentoring of S.B.O. and members of the Blokesch and Bitbol laboratory for fruitful discussions. This work was supported by grants from the Swiss National Science Foundation (310030_185022) and the European Research Council (724630) and an International Research Scholarship by the Howard Hughes Medical Institute awarded to M.B. (55008726). R. S. and A.-F. B. acknowledge funding from the European Research Council (ERC) under the European Union’s Horizon 2020 research and innovation programme (grant agreement No. 851173, to A.-F. B.).

## Author contributions

S.B.O and M.B. conceived the project, designed and analysed the biological experiments. S.B.O. constructed strains and plasmids and performed all biological experiments. R.S. and A.-F.B. designed and analysed the computational model and the numerical simulations. R.S. performed all numerical simulations. A.L. performed bioinformatic analyses. S.B.O. and M.B. wrote the manuscript with input from A.L., R.S., and A.-F.B. M.B. and A.-F.B. acquired funding and supervised the project.

## Supplementary Figure legends

**Fig. S1. Refinement of Bacterial Competition Assays. A)** Relative fitness of antibiotic/fluorescent marker-carrying strains used for the construction of predator and prey strains, competed against the control strain (A1552Δ*lacZ*). **B)** LB medium batch-to-batch variability in the aggregation level can be counteracted by the supplementation of divalent cations. The aggregation level is determined by the ratio of the culture’s OD_600_ pre/post vortex. The horizontal dotted line represents the ratio around which no aggregation occurs. All presented values are the mean of 3 repeats, with error bars indicating the standard deviation. Significant differences were determined by a two-tailed Student’s t-test.

**Fig. S2. Evolution over time of a simulated aggregate with matching T4P and T6SS killing.** The simulation is performed as described in the methods section, where all parameter values are given. Prey and predators are represented by green and red markers, respectively. Lysing cells are represented by black markers. Within each panel, the number of bacteria forming the aggregate of interest is indicated, and the black scale bar shows the length of 5 marker diameters. In the second panel (1 minute), we show two randomly chosen small aggregates (the dashed line separating the two plots indicates that they are not close to each other), but note that other aggregates or isolated bacteria later end up within the aggregate of interest. In the first panel (0 minute), we display the content of an arbitrary cube, of edge about 23 cell diameters, extracted from the system.

**Fig. S3. Comparison of Large Cluster-encoded Immunity Proteins.** The heat map displays the fraction identity (fraction of identical residues) among core immunity proteins encoded in the T6SS large cluster (LC) from patient and environmental isolates used in this study, along with strain representatives of known families [17,61]. Green indicates a fraction identity of 1 (same family and subfamily), yellow indicates a fraction identity of >0.3 (same family), and red indicates a fraction identity of ≤0.3 (different family). Families are labelled by letters on the side of the heatmap, and stars denote novel families. After manual inspection, SL6Y was classified into its own family as the identity values corresponding to family H were at the threshold level and did not encompass all of the family H strains.

**Fig. S4. Comparison of Auxiliary Cluster 1-encoded Immunity Proteins.** The heat map illustrates the fraction identity (fraction of identical residues) among core immunity proteins located in T6SS auxiliary cluster 1 (Aux1) from patient and environmental isolates used in this study, along with strain representatives of known families [17,61]. Green indicates a fraction identity of 1 (same family and subfamily), yellow indicates a fraction identity of >0.3 (same family), and red indicates a fraction identity of ≤0.3 (different family). Families are labelled by letters on the side of the heatmap.

**Fig. S5. Comparison of Auxiliary Cluster 2-encoded Immunity Proteins.** The heat map illustrates the fraction identity (fraction of identical residues) among core immunity proteins located in T6SS auxiliary cluster 2 (Aux2) from patient and environmental isolates used in this study, along with strain representatives of known families [17,61]. Green indicates a fraction identity of 1 (same family and subfamily), yellow indicates a fraction identity of >0.3 (same family), and red indicates a fraction identity of ≤0.3 (different family). Families are labelled by letters on the side of the heatmap.

**Fig. S6. Comparison of Auxiliary Cluster 3-encoded Immunity Proteins.** The heat map illustrates the fraction identity (fraction of identical residues) among core immunity proteins located in T6SS auxiliary cluster 3 (Aux3) from patient and environmental isolates used in this study. Green indicates a fraction identity of 1 (same family and subfamily), yellow indicates a fraction identity of >0.3 (same family), and red indicates a fraction identity of ≤0.3 (different family). Families are labelled by letters on the side of the heatmap. Dashed lines indicate a longer predicted sequence at the beginning of the proteins.

**Fig. S7. Comparison of Auxiliary Cluster 4-encoded Immunity Proteins.** The heat map illustrates the fraction identity (fraction of identical residues) among core immunity proteins located in T6SS auxiliary cluster 4 (Aux4) from patient and environmental isolates used in this study. Green indicates a fraction identity of 1 (same family and subfamily), yellow indicates a fraction identity of >0.3 (same family), and red indicates a fraction identity of ≤0.3 (different family). Families are labelled by letters on the side of the heatmap.

**Fig. S8. Comparison of Auxiliary Cluster 5-encoded Immunity Proteins.** The heat map illustrates the fraction identity (fraction of identical residues) among core immunity proteins located in T6SS auxiliary cluster 5 (Aux5) from patient and environmental isolates used in this study. Green indicates a fraction identity of 1 (same family and subfamily), yellow indicates a fraction identity of >0.3 (same family), and red indicates a fraction identity of ≤0.3 (different family). Families are labelled by letters on the side of the heatmap.

**Fig. S9. Comparison of PilA proteins.** The heat map displaying fraction identity (fraction of identical residues) of PilA from patient and environmental isolates used in this study, which includes A1552 as a pandemic representative and *V. mimicus* (ATCC33655) as the outgroup. Green indicates fraction identity of 1, yellow indicates fraction identity >0.75, red fraction identity >0.5, and dark red fraction identity ≤0.5. The experimentally proven ability of the PilA variants to self-interact is indicated above the heatmap.

**Fig. S10. The *pilA* gene likely moves by horizontal gene transfer.** The figure compared the core gene-based cladogram of the 39 *V. cholerae* strains studied (on the left side) with the *pilA* nucleotide sequences-based cladogram (on the right side). Coloured boxes highlight supported clades of relatively related strains, which are also represented by coloured circles in the *pilA* cladogram. The incongruence between the genome-based and *pilA* reconstructions suggests horizontal gene transfer of *pilA*. *V. mimicus* (ATCC33655) is used as an outgroup. Statistical significance was verified using 100 bootstraps, and nodes with bootstrap values below 60 are collapsed.

**Fig. S11. Pilus diversity provides T6SS protection by spatial segregation.** Microscopy images of co-culture experiments between T6SS-competent (Parent) or non-functional (Δ*vasK*) predator strains, and T6SS-sensitive (Δ4E/I) prey strains. Predator and prey strains are either carrying matching (upper panels), or diverse (lower panels) PilA variants. Phase contrast, a merge of prey (sfGFP, in green) and predator strains (mCherry, in red), and SYTOX Blue dead cell stain (blue) channels are displayed. Scalebar indicates 25 µm.

**Fig. S12. Microscopy of strain pairs with promiscuous T4P combinations.** Images of co-culture experiments between T6SS-competent (predator) and T6SS-sensitive (prey) strains, displaying all PilA variants, known to exhibit levels of promiscuousness. Phase contrast, a merge of prey (sfGFP, green) and predator strains (mCherry, red), and SYTOX Blue dead cell stain (blue) channels are displayed. Scalebar indicates 25 µm.

**Fig. S13. 2D simulations yield a similar phenomenology as 3D simulations.** In addition to the 3D simulations (see methods section), 2D simulations were performed in a triangular lattice, with the same division rate, diffusion coefficient, binding energies, total firing rate, and lysis rate as in the 3D case. The 5 panels show zooms from snapshots taken after 1 h of 2D simulations where 100×100 triangular lattices were initialized with 50 prey (green markers) and 50 predators (red markers) placed uniformly at random (note that this yields a larger density than in our 3D simulations). Lysing cells are represented by black markers. The black scale bar indicates the length of 5 marker diameters. On the left: no interaction between prey and predators (*E_cross_* = 0). The top panel illustrates the complete absence of T4P, while the bottom panel corresponds to the non-matching T4P case. On the right: prey and predators interact via T4P with a non-zero binding energy (*E_cross_* > 0). The top panel shows the matching T4P case, and the middle and bottom panels correspond to promiscuous T4P, with *E_cross_* equal to 1.5*kT* and 1*kT*, respectively.

**Fig. S14. A similar phenomenology is obtained by four variants of the model.** On the left are 2D simulations, and on the right are 3D simulations. The top panels illustrate simulations that include the diffusion of aggregates as a whole, while the bottom panels depict simulations conducted without such diffusion (see methods). All panels show zooms from snapshots taken after 1 h of simulations performed as described in the methods section for 3D simulations, and as described in the legend of Figure S13 for 2D simulations. Prey and predators are represented by green and red markers, respectively, while lysing cells are represented by black markers. The black scale bar indicates the length of 5 marker diameters. The main behaviours (aggregation or no aggregation) remain consistent across all variants of the model. Notably, when aggregates do not diffuse, multiple prey and predator aggregates are formed after one hour. In contrast, when they diffuse, aggregates of a particular type coalesce, resulting in a single aggregate of prey and a single aggregate of predators.

**Fig. S15. Aggregation is facilitated by T4P with sufficient binding energy.** The plot displays the mean local density, which represents the average fraction of occupied neighbouring sites of a bacterium, plotted against the T4P binding energy *E* at steady state. In this figure, we consider a single type of bacteria (either prey or predators) diffusing and interacting through T4P with a specific attractive energy *E*, but without division or killing. The system can then be mapped to a lattice fluid model, and a Monte Carlo simulation of its steady state was performed. The blue curve represents the 3D model where bacteria diffuse on a 40 × 40 × 40 body-centred cubic lattice, and the yellow curve corresponds to the 2D model where bacteria diffuse on a 100 × 100 triangular lattice. In both cases, the system comprises 100 bacteria.

## Supplementary Tables

**Table S1. Bacterial strains and plasmids used in this study.**

**Table S2. Information on genomes used for bioinformatical analysis.**

## Supplementary Movies

**Movie S1. Simulation with matching T4P; view of a fixed cube.** This video shows the evolution of an aggregate over one hour in the case of matching T4P with T6SS killing enabled. The region of interest is a cube extracted from the full simulation volume, whose position remains fixed throughout the video. Prey and predators are represented by green and red markers, respectively. Lysing cells are represented by black markers. The playback speed of the video compared to real time is indicated at the top right of the video.

**Movie S2. Simulation without T4P; view of a fixed cube.** This video shows the evolution of an aggregate over one hour in the case without T4P but with T6SS killing enabled. The region of interest is a cube extracted from the full simulation volume, whose position remains fixed throughout the video. Prey and predators are represented by green and red markers, respectively. Lysing cells are represented by black markers. The playback speed of the video compared to real time is indicated at the top right of the video.

**Movie S3. Simulation with matching T4P but no T6SS.** This video shows the evolution of an aggregate over one hour in the case of matching T4P with T6SS killing disabled. The region of interest is a cube extracted from the full simulation volume, centred on an aggregate of interest. However, note that at the beginning of the video, when aggregates are just starting to form, the aforementioned cube is not centred on any aggregate yet. Prey and predators are represented by green and red markers, respectively. Lysing cells are represented by black markers. The playback speed of the video compared to real time is indicated at the top right of the video.

**Movie S4. Simulation with matching T4P.** This video shows the evolution of an aggregate over one hour in the case of matching T4P with T6SS killing enabled. The region of interest is a cube extracted from the full simulation volume, centred on an aggregate of interest. However, note that at the beginning of the video, when aggregates are just starting to form, the aforementioned cube is not centred on any aggregate yet. Prey and predators are represented by green and red markers, respectively. Lysing cells are represented by black markers. The playback speed of the video compared to real time is indicated at the top right of the video.

**Movie S5. Simulation with non-matching T4P.**This video shows the evolution of an aggregate over one hour in the case of non-matching T4P with T6SS killing enabled. The region of interest is a cube extracted from the full simulation volume, centred on an aggregate of interest. However, note that at the beginning of the video, when aggregates are just starting to form, the aforementioned cube is not centred on any aggregate yet. Prey and predators are represented by green and red markers, respectively. Lysing cells are represented by black markers. The playback speed of the video compared to real time is indicated at the top right of the video.

**Movie S6. Simulation with promiscuous T4P.** This video shows the evolution of an aggregate over one hour in the case of promiscuous T4P (*E_cross_* = 1*kT*) with T6SS killing enabled. The region of interest is a cube extracted from the full simulation volume, centred on an aggregate of interest. However, note that at the beginning of the video, when aggregates are just starting to form, the aforementioned cube is not centred on any aggregate yet. Prey and predators are represented by green and red markers, respectively. Lysing cells are represented by black markers. The playback speed of the video compared to real time is indicated at the top right of the video.

