## Supplementary figures for "Interactions between pili affect the outcome of bacterial competition driven by the type VI secretion system"

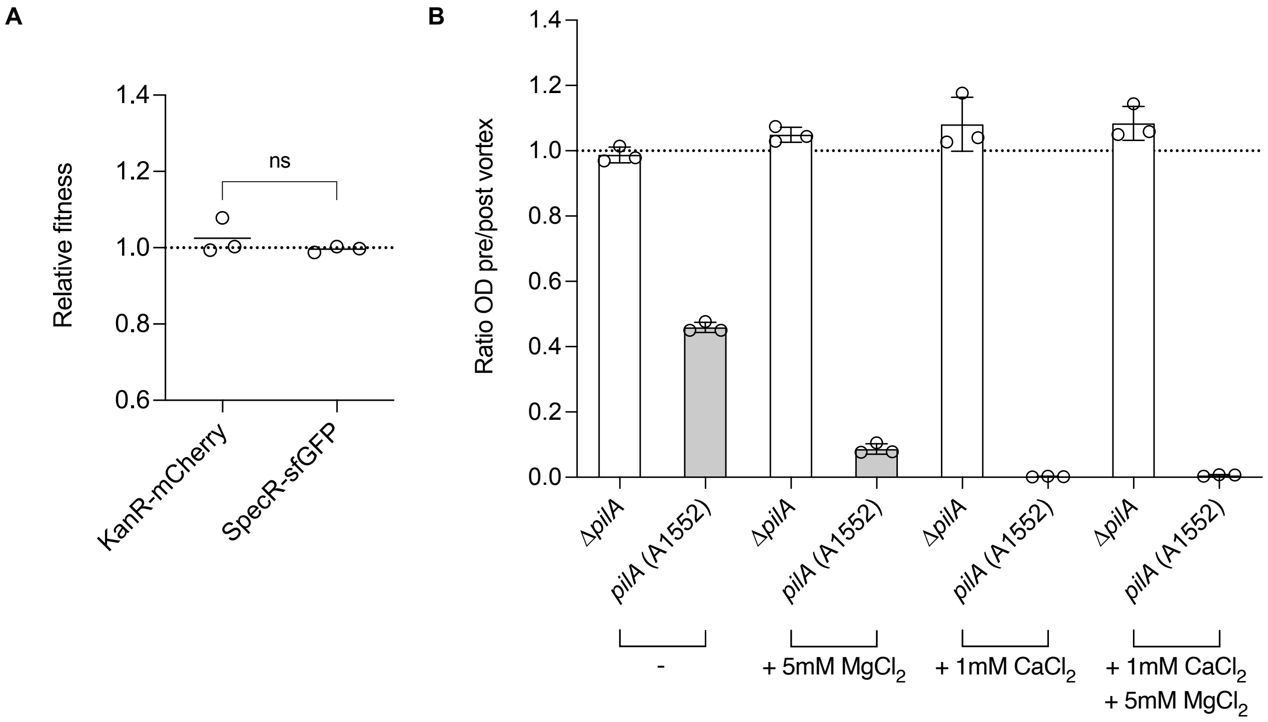


**Fig. S1. Refinement of Bacterial Competition Assays. A)** Relative fitness of antibiotic/fluorescent marker-carrying strains used for the construction of predator and prey strains, competed against the control strain (A1552Δ*lacZ*). **B)** LB medium batch-to-batch variability in the aggregation level can be counteracted by the supplementation of divalent cations. The aggregation level is determined by the ratio of the culture's $OD_{600}$ pre/post vortex. The horizontal dotted line represents the ratio around which no aggregation occurs. All presented values are the mean of 3 repeats, with error bars indicating the standard deviation. Significant differences were determined by a two-tailed Student's t-test.

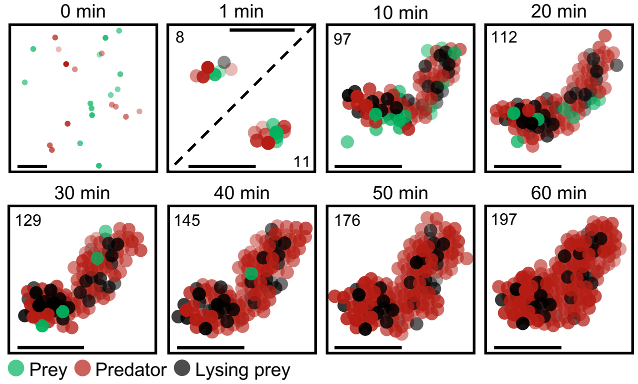


**Fig. S2. Evolution over time of a simulated aggregate with matching T4P and T6SS killing.** The simulation is performed as described in the methods section, where all parameter values are given. Prey and predators are represented by green and red markers, respectively. Lysing cells are represented by black markers. Within each panel, the number of bacteria forming the aggregate of interest is indicated, and the black scale bar shows the length of 5 marker diameters. In the second panel (1 minute), we show two randomly chosen small aggregates (the dashed line separating the two plots indicates that they are not close to each other), but note that other aggregates or isolated bacteria later end up within the aggregate of interest. In the first panel (0 minute), we display the content of an arbitrary cube, of edge about 23 cell diameters, extracted from the system.

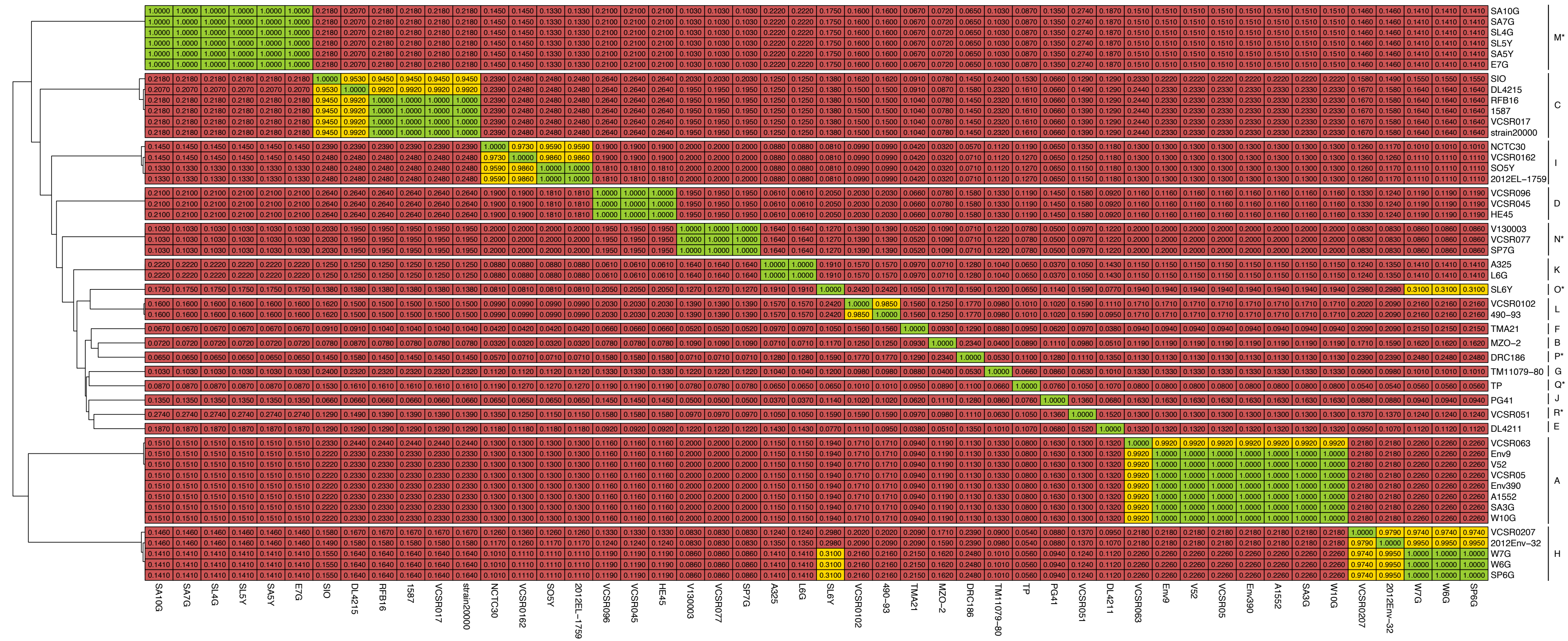


**Fig. S3. Comparison of Large Cluster-encoded Immunity Proteins.** The heat map displays the fraction identity (fraction of identical residues) among core immunity proteins encoded in the T6SS large cluster (LC) from patient and environmental isolates used in this study, along with strain representatives of known families [17,61]. Green indicates a fraction identity of 1 (same family and subfamily), yellow indicates a fraction identity of >0.3 (same family), and red indicates a fraction identity of ≤0.3 (different family). Families are labelled by letters on the side of the heatmap, and stars denote novel families. After manual inspection, SL6Y was classified into its own family as the identity values corresponding to family H were at the threshold level and did not encompass all of the family H strains.

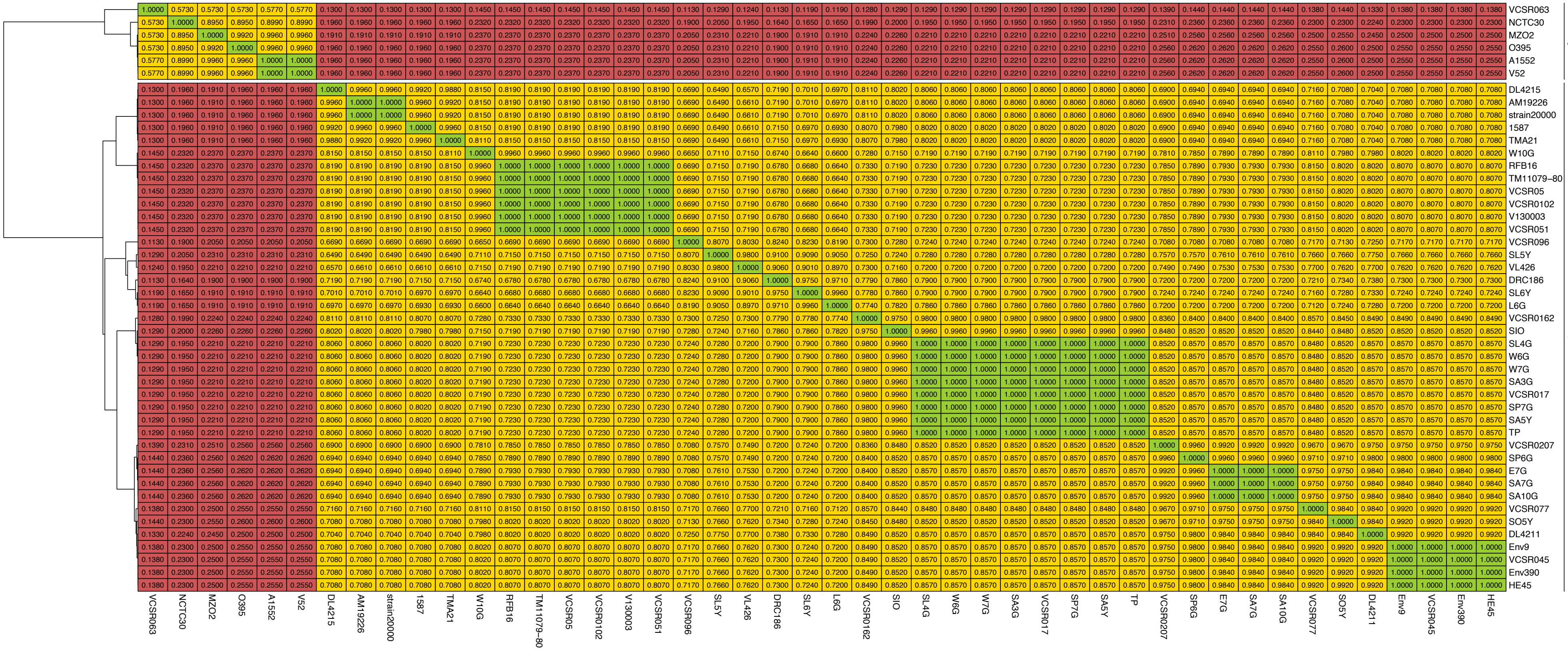


**Fig. S4. Comparison of Auxiliary Cluster 1-encoded Immunity Proteins.** The heat map illustrates the fraction identity (fraction of identical residues) among core immunity proteins located in T6SS auxiliary cluster 1 (Aux1) from patient and environmental isolates used in this study, along with strain representatives of known families [17,61]. Green indicates a fraction identity of 1 (same family and subfamily), yellow indicates a fraction identity of >0.3 (same family), and red indicates a fraction identity of ≤0.3 (different family). Families are labelled by letters on the side of the heatmap.

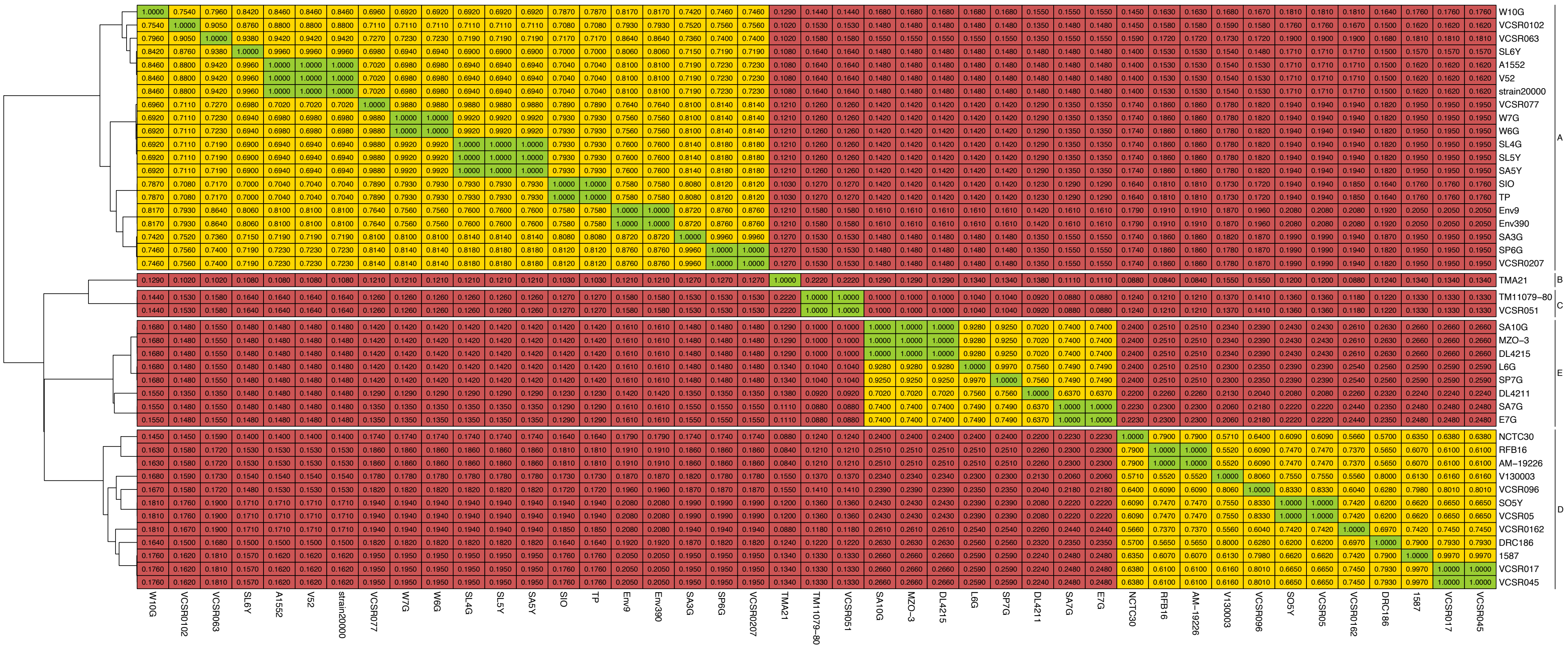


**Fig. S5. Comparison of Auxiliary Cluster 2-encoded Immunity Proteins.** The heat map illustrates the fraction identity (fraction of identical residues) among core immunity proteins located in T6SS auxiliary cluster 2 (Aux2) from patient and environmental isolates used in this study, along with strain representatives of known families [17,61]. Green indicates a fraction identity of 1 (same family and subfamily), yellow indicates a fraction identity of >0.3 (same family), and red indicates a fraction identity of ≤0.3 (different family). Families are labelled by letters on the side of the heatmap.

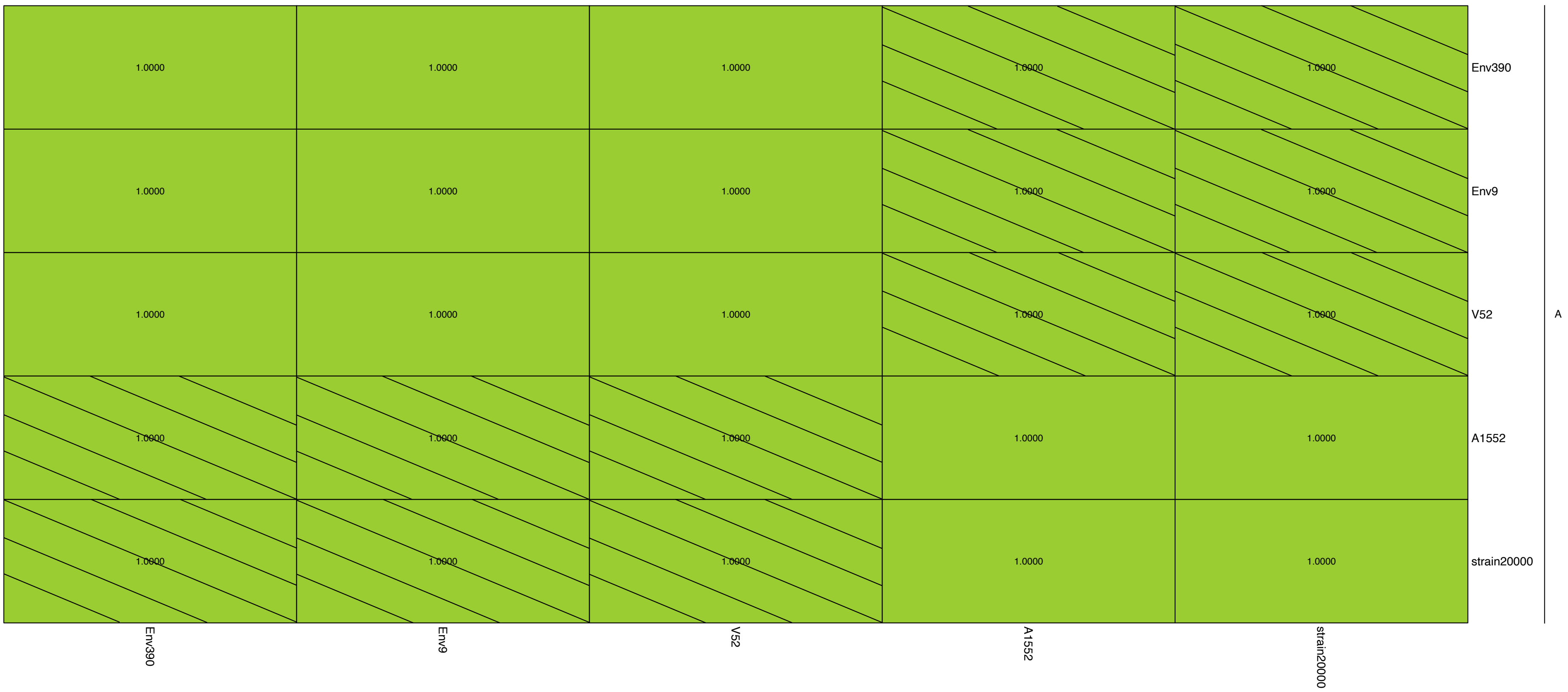


**Fig. S6. Comparison of Auxiliary Cluster 3-encoded Immunity Proteins.** The heat map illustrates the fraction identity (fraction of identical residues) among core immunity proteins located in T6SS auxiliary cluster 3 (Aux3) from patient and environmental isolates used in this study. Green indicates a fraction identity of 1 (same family and subfamily), yellow indicates a fraction identity of >0.3 (same family), and red indicates a fraction identity of ≤0.3 (different family). Families are labelled by letters on the side of the heatmap. Dashed lines indicate a longer predicted sequence at the beginning of the proteins.

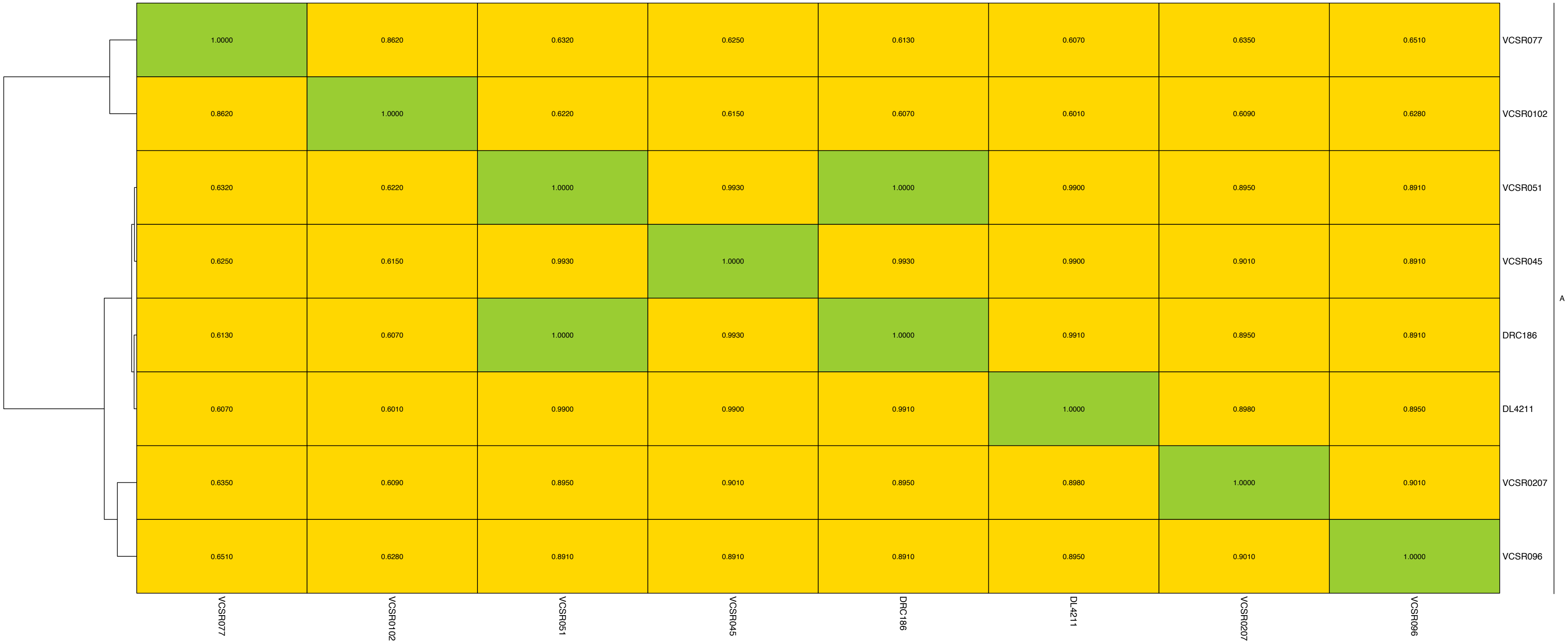


**Fig. S7. Comparison of Auxiliary Cluster 4-encoded Immunity Proteins.** The heat map illustrates the fraction identity (fraction of identical residues) among core immunity proteins located in T6SS auxiliary cluster 4 (Aux4) from patient and environmental isolates used in this study. Green indicates a fraction identity of 1 (same family and subfamily), yellow indicates a fraction identity of >0.3 (same family), and red indicates a fraction identity of ≤0.3 (different family). Families are labelled by letters on the side of the heatmap.

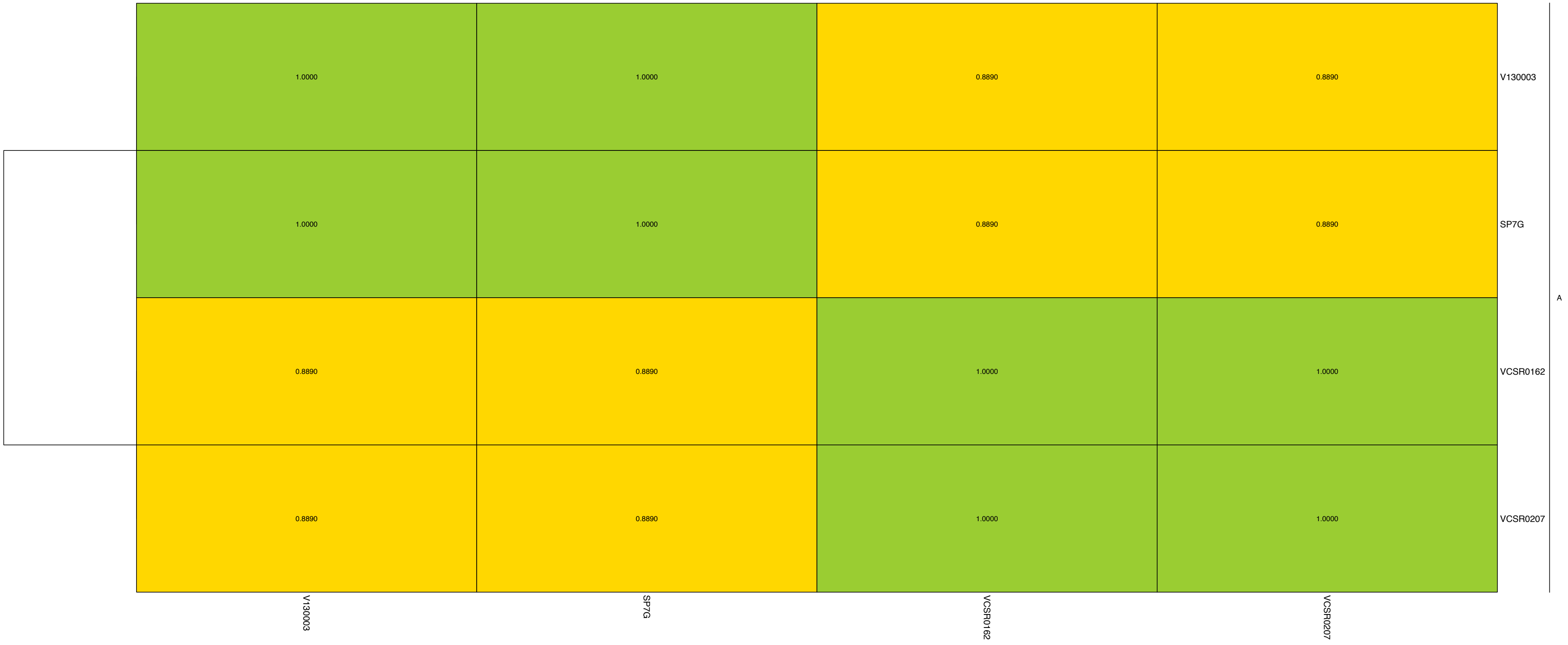


**Fig. S8. Comparison of Auxiliary Cluster 5-encoded Immunity Proteins.** The heat map illustrates the fraction identity (fraction of identical residues) among core immunity proteins located in T6SS auxiliary cluster 5 (Aux5) from patient and environmental isolates used in this study. Green indicates a fraction identity of 1 (same family and subfamily), yellow indicates a fraction identity of >0.3 (same family), and red indicates a fraction identity of ≤0.3 (different family). Families are labelled by letters on the side of the heatmap.

| | Outgroup | Self-interacting | | | | | | | | Intermediate | | Non-interacting | | | | | | | Unknown | |
|---|---|---|---|---|---|---|---|---|---|---|---|---|---|---|---|---|---|---|---|---|
| | *V. mimicus* | A1552 | SP6G | SP7G | L6G | SA3G | TP | DL4215 | 1587 | SL6Y | DRC186 | SA5Y | V52 | W6G | SA7G | SO5Y | SIO | W10G | SL5Y | SL4G |
| *V. mimicus* | 1.0000 | 0.1540 | 0.1790 | 0.1430 | 0.1300 | 0.2050 | 0.1690 | 0.1950 | 0.1840 | 0.1950 | 0.1410 | 0.2310 | 0.1690 | 0.1670 | 0.1430 | 0.1670 | 0.1790 | 0.1710 | 0.1950 | 0.2310 |
| A1552 | 0.1540 | 1.0000 | 0.5560 | 0.3740 | 0.3710 | 0.5400 | 0.5450 | 0.5440 | 0.4630 | 0.4960 | 0.4130 | 0.5660 | 0.5180 | 0.4170 | 0.4930 | 0.4170 | 0.4410 | 0.4890 | 0.4960 | 0.5660 |
| SP6G | 0.1790 | 0.5560 | 1.0000 | 0.3360 | 0.3410 | 0.6010 | 0.5070 | 0.6480 | 0.5210 | 0.5110 | 0.4410 | 0.6200 | 0.5190 | 0.4860 | 0.4710 | 0.4860 | 0.5160 | 0.5220 | 0.5110 | 0.6200 |
| SP7G | 0.1430 | 0.3740 | 0.3360 | 1.0000 | 0.4640 | 0.3750 | 0.4000 | 0.3660 | 0.3360 | 0.4560 | 0.3890 | 0.3830 | 0.3630 | 0.3540 | 0.3720 | 0.3540 | 0.3870 | 0.3650 | 0.4560 | 0.3830 |
| L6G | 0.1300 | 0.3710 | 0.3410 | 0.4640 | 1.0000 | 0.3330 | 0.4170 | 0.3510 | 0.3360 | 0.3920 | 0.3460 | 0.3440 | 0.3360 | 0.3280 | 0.3410 | 0.3280 | 0.3390 | 0.3570 | 0.3920 | 0.3440 |
| SA3G | 0.2050 | 0.5400 | 0.6010 | 0.3750 | 0.3330 | 1.0000 | 0.5190 | 0.5460 | 0.4960 | 0.5150 | 0.4750 | 0.7540 | 0.5300 | 0.4890 | 0.4890 | 0.4890 | 0.4800 | 0.4810 | 0.5150 | 0.7540 |
| TP | 0.1690 | 0.5450 | 0.5070 | 0.4000 | 0.4170 | 0.5190 | 1.0000 | 0.4590 | 0.3970 | 0.5940 | 0.4180 | 0.5220 | 0.5320 | 0.4690 | 0.4450 | 0.4690 | 0.4470 | 0.4520 | 0.5940 | 0.5220 |
| DL4215 | 0.1950 | 0.5440 | 0.6480 | 0.3660 | 0.3510 | 0.5460 | 0.4590 | 1.0000 | 0.4680 | 0.5040 | 0.3880 | 0.5670 | 0.4730 | 0.4960 | 0.4490 | 0.4960 | 0.5390 | 0.5040 | 0.5040 | 0.5670 |
| 1587 | 0.1840 | 0.4630 | 0.5210 | 0.3360 | 0.3360 | 0.4960 | 0.3970 | 0.4680 | 1.0000 | 0.4540 | 0.4370 | 0.4740 | 0.5450 | 0.5070 | 0.4810 | 0.5070 | 0.4720 | 0.5040 | 0.4540 | 0.4740 |
| SL6Y | 0.1950 | 0.4960 | 0.5110 | 0.4560 | 0.3920 | 0.5150 | 0.5940 | 0.5040 | 0.4540 | 1.0000 | 0.4140 | 0.5080 | 0.4690 | 0.4620 | 0.4440 | 0.4620 | 0.4840 | 0.4610 | 1.0000 | 0.5080 |
| DRC186 | 0.1410 | 0.4130 | 0.4410 | 0.3890 | 0.3460 | 0.4750 | 0.4180 | 0.3880 | 0.4370 | 0.4140 | 1.0000 | 0.4500 | 0.4320 | 0.4710 | 0.5070 | 0.4710 | 0.5240 | 0.5380 | 0.4140 | 0.4500 |
| SA5Y | 0.2310 | 0.5660 | 0.6200 | 0.3830 | 0.3440 | 0.7540 | 0.5220 | 0.5670 | 0.4740 | 0.5080 | 0.4500 | 1.0000 | 0.5380 | 0.4810 | 0.4890 | 0.4810 | 0.5120 | 0.5040 | 0.5080 | 1.0000 |
| V52 | 0.1690 | 0.5180 | 0.5190 | 0.3630 | 0.3360 | 0.5300 | 0.5320 | 0.4730 | 0.5450 | 0.4690 | 0.4320 | 0.5380 | 1.0000 | 0.4970 | 0.4810 | 0.5070 | 0.4650 | 0.5310 | 0.4690 | 0.5380 |
| W6G | 0.1670 | 0.4170 | 0.4860 | 0.3540 | 0.3280 | 0.4890 | 0.4690 | 0.4960 | 0.5070 | 0.4620 | 0.4710 | 0.4810 | 0.4970 | 1.0000 | 0.5830 | 1.0000 | 0.7330 | 0.5410 | 0.4620 | 0.4810 |
| SA7G | 0.1430 | 0.4930 | 0.4710 | 0.3720 | 0.3410 | 0.4890 | 0.4450 | 0.4490 | 0.4810 | 0.4440 | 0.5070 | 0.4890 | 0.4810 | 0.5830 | 1.0000 | 0.5830 | 0.6080 | 0.5480 | 0.4440 | 0.4890 |
| SO5Y | 0.1670 | 0.4170 | 0.4860 | 0.3540 | 0.3280 | 0.4890 | 0.4690 | 0.4960 | 0.5070 | 0.4620 | 0.4710 | 0.4810 | 0.5070 | 1.0000 | 0.5830 | 1.0000 | 0.7330 | 0.5410 | 0.4620 | 0.4810 |
| SIO | 0.1790 | 0.4410 | 0.5160 | 0.3870 | 0.3390 | 0.4800 | 0.4470 | 0.5390 | 0.4720 | 0.4840 | 0.5240 | 0.5120 | 0.4650 | 0.7330 | 0.6080 | 0.7330 | 1.0000 | 0.5890 | 0.4840 | 0.5120 |
| W10G | 0.1710 | 0.4890 | 0.5220 | 0.3650 | 0.3570 | 0.4810 | 0.4520 | 0.5040 | 0.5040 | 0.4610 | 0.5380 | 0.5040 | 0.5310 | 0.5410 | 0.5480 | 0.5410 | 0.5890 | 1.0000 | 0.4610 | 0.5040 |
| SL5Y | 0.1950 | 0.4960 | 0.5110 | 0.4560 | 0.3920 | 0.5150 | 0.5940 | 0.5040 | 0.4540 | 1.0000 | 0.4140 | 0.5080 | 0.4690 | 0.4620 | 0.4440 | 0.4620 | 0.4840 | 0.4610 | 1.0000 | 0.5080 |
| SL4G | 0.2310 | 0.5660 | 0.6200 | 0.3830 | 0.3440 | 0.7540 | 0.5220 | 0.5670 | 0.4740 | 0.5080 | 0.4500 | 1.0000 | 0.5380 | 0.4810 | 0.4890 | 0.4810 | 0.5120 | 0.5040 | 0.5080 | 1.0000 |
| W7G | 0.1670 | 0.4170 | 0.4860 | 0.3540 | 0.3280 | 0.4890 | 0.4690 | 0.4960 | 0.5070 | 0.4620 | 0.4710 | 0.4810 | 0.4970 | 1.0000 | 0.5830 | 1.0000 | 0.7330 | 0.5410 | 0.4620 | 0.4810 |
| SA10G | 0.1430 | 0.4930 | 0.4710 | 0.3720 | 0.3410 | 0.4890 | 0.4450 | 0.4490 | 0.4810 | 0.4440 | 0.5070 | 0.4890 | 0.4810 | 0.5830 | 1.0000 | 0.5830 | 0.6080 | 0.5480 | 0.4440 | 0.4890 |
| E7G | 0.1430 | 0.4930 | 0.4710 | 0.3720 | 0.3410 | 0.4890 | 0.4450 | 0.4490 | 0.4810 | 0.4440 | 0.5070 | 0.4890 | 0.4810 | 0.5830 | 1.0000 | 0.5830 | 0.6080 | 0.5480 | 0.4440 | 0.4890 |
| DL4211 | 0.1950 | 0.4960 | 0.5110 | 0.4560 | 0.3920 | 0.5150 | 0.5940 | 0.5040 | 0.4540 | 1.0000 | 0.4140 | 0.5080 | 0.4690 | 0.4620 | 0.4440 | 0.4620 | 0.4840 | 0.4610 | 1.0000 | 0.5080 |
| VCSR05 | 0.2210 | 0.5190 | 0.5260 | 0.4400 | 0.4000 | 0.5450 | 0.5780 | 0.4740 | 0.4310 | 0.7990 | 0.4360 | 0.5000 | 0.4730 | 0.4550 | 0.4810 | 0.4550 | 0.4840 | 0.4770 | 0.7990 | 0.5000 |
| VCSR0207 | 0.1970 | 0.5390 | 0.5530 | 0.4270 | 0.3870 | 0.5110 | 0.5350 | 0.5080 | 0.4230 | 0.7370 | 0.4320 | 0.5190 | 0.5000 | 0.4500 | 0.4700 | 0.4500 | 0.4800 | 0.4920 | 0.7370 | 0.5190 |
| VCSR096 | 0.1950 | 0.5370 | 0.4700 | 0.4170 | 0.4250 | 0.5110 | 0.9050 | 0.4440 | 0.3970 | 0.5940 | 0.4180 | 0.5220 | 0.5240 | 0.4450 | 0.4220 | 0.4450 | 0.4390 | 0.4370 | 0.5940 | 0.5220 |
| V130003 | 0.1950 | 0.5370 | 0.4700 | 0.4170 | 0.4250 | 0.5110 | 0.9050 | 0.4440 | 0.3970 | 0.5940 | 0.4180 | 0.5220 | 0.5240 | 0.4450 | 0.4220 | 0.4450 | 0.4390 | 0.4370 | 0.5940 | 0.5220 |
| Env390 | 0.1540 | 1.0000 | 0.5560 | 0.3740 | 0.3710 | 0.5400 | 0.5450 | 0.5440 | 0.4630 | 0.4960 | 0.4130 | 0.5660 | 0.5180 | 0.4170 | 0.4930 | 0.4170 | 0.4410 | 0.4890 | 0.4960 | 0.5660 |
| Env9 | 0.1540 | 1.0000 | 0.5560 | 0.3740 | 0.3710 | 0.5400 | 0.5450 | 0.5440 | 0.4630 | 0.4960 | 0.4130 | 0.5660 | 0.5180 | 0.4170 | 0.4930 | 0.4170 | 0.4410 | 0.4890 | 0.4960 | 0.5660 |
| VCSR063 | 0.1690 | 0.5260 | 0.5260 | 0.3550 | 0.3280 | 0.5230 | 0.5240 | 0.4810 | 0.5380 | 0.4620 | 0.4240 | 0.5460 | 0.9940 | 0.4900 | 0.4740 | 0.5000 | 0.4720 | 0.5380 | 0.4620 | 0.5460 |
| VCSR051 | 0.1430 | 0.4350 | 0.4710 | 0.3510 | 0.3480 | 0.4850 | 0.4490 | 0.4560 | 0.4810 | 0.4200 | 0.5110 | 0.4930 | 0.4110 | 0.4830 | 0.5580 | 0.4830 | 0.5280 | 0.5470 | 0.4200 | 0.4930 |
| VCSR0102 | 0.1430 | 0.4350 | 0.4710 | 0.3510 | 0.3480 | 0.4850 | 0.4490 | 0.4560 | 0.4810 | 0.4200 | 0.5110 | 0.4930 | 0.4050 | 0.4770 | 0.5580 | 0.4770 | 0.5280 | 0.5470 | 0.4200 | 0.4930 |
| VCSR0162 | 0.1690 | 0.4670 | 0.4710 | 0.3620 | 0.3510 | 0.4670 | 0.4410 | 0.4640 | 0.5260 | 0.4360 | 0.5220 | 0.5040 | 0.4850 | 0.5360 | 0.5430 | 0.5360 | 0.5690 | 0.5750 | 0.4360 | 0.5040 |
| RFB16 | 0.1710 | 0.4550 | 0.4590 | 0.3620 | 0.3620 | 0.4770 | 0.4170 | 0.4550 | 0.5220 | 0.4110 | 0.5570 | 0.4700 | 0.4810 | 0.5520 | 0.5510 | 0.5520 | 0.5600 | 0.8140 | 0.4110 | 0.4700 |
| VCSR017 | 0.1450 | 0.4560 | 0.4750 | 0.3410 | 0.3250 | 0.4780 | 0.4200 | 0.4110 | 0.5960 | 0.4200 | 0.4810 | 0.4400 | 0.4920 | 0.4530 | 0.5180 | 0.4530 | 0.4880 | 0.4960 | 0.4200 | 0.4400 |
| NCTC30 | 0.1580 | 0.4530 | 0.4570 | 0.3250 | 0.3170 | 0.4220 | 0.3710 | 0.4230 | 0.6200 | 0.4050 | 0.4560 | 0.4300 | 0.4850 | 0.4450 | 0.4890 | 0.4450 | 0.4800 | 0.5110 | 0.4050 | 0.4300 |
| VCSR077 | 0.1300 | 0.3920 | 0.3360 | 0.4510 | 0.5930 | 0.3770 | 0.4390 | 0.3480 | 0.3660 | 0.4340 | 0.3620 | 0.3720 | 0.3770 | 0.3810 | 0.3700 | 0.3810 | 0.3920 | 0.3740 | 0.4340 | 0.3720 |
| strain20000 | 0.1430 | 0.3980 | 0.3610 | 0.4660 | 0.5680 | 0.4200 | 0.4210 | 0.3480 | 0.3660 | 0.4050 | 0.3750 | 0.3920 | 0.3640 | 0.3510 | 0.3410 | 0.3510 | 0.3610 | 0.3820 | 0.4050 | 0.3920 |
| VCSR045 | 0.1430 | 0.3740 | 0.3360 | 0.9430 | 0.4500 | 0.3830 | 0.4170 | 0.3590 | 0.3440 | 0.4480 | 0.4130 | 0.3750 | 0.3470 | 0.3700 | 0.3880 | 0.3700 | 0.3870 | 0.3650 | 0.4480 | 0.3750 |

| | Unknown | | | | | | | | | | | | | | | | | | | |
|---|---|---|---|---|---|---|---|---|---|---|---|---|---|---|---|---|---|---|---|---|
| | W7G | SA10G | E7G | DL4211 | VCSR05 | VCSR0207 | VCSR096 | V130003 | Env390 | Env9 | VCSR063 | VCSR051 | VCSR0102 | VCSR0162 | RFB16 | VCSR017 | NCTC30 | VCSR077 | strain20000 | VCSR045 |
| *V. mimicus* | 0.1670 | 0.1430 | 0.1430 | 0.1950 | 0.2210 | 0.1970 | 0.1950 | 0.1950 | 0.1540 | 0.1540 | 0.1690 | 0.1430 | 0.1430 | 0.1690 | 0.1710 | 0.1450 | 0.1580 | 0.1300 | 0.1430 | 0.1430 |
| A1552 | 0.4170 | 0.4930 | 0.4930 | 0.4960 | 0.5190 | 0.5390 | 0.5370 | 0.5370 | 1.0000 | 1.0000 | 0.5260 | 0.4350 | 0.4350 | 0.4670 | 0.4550 | 0.4560 | 0.4530 | 0.3920 | 0.3980 | 0.3740 |
| SP6G | 0.4860 | 0.4710 | 0.4710 | 0.5110 | 0.5260 | 0.5530 | 0.4700 | 0.4700 | 0.5560 | 0.5560 | 0.5260 | 0.4710 | 0.4710 | 0.4710 | 0.4590 | 0.4750 | 0.4570 | 0.3360 | 0.3610 | 0.3360 |
| SP7G | 0.3540 | 0.3720 | 0.3720 | 0.4560 | 0.4400 | 0.4270 | 0.4170 | 0.4170 | 0.3740 | 0.3740 | 0.3550 | 0.3510 | 0.3510 | 0.3620 | 0.3620 | 0.3410 | 0.3250 | 0.4510 | 0.4660 | 0.9430 |
| L6G | 0.3280 | 0.3410 | 0.3410 | 0.3920 | 0.4000 | 0.3870 | 0.4250 | 0.4250 | 0.3710 | 0.3710 | 0.3280 | 0.3480 | 0.3480 | 0.3510 | 0.3620 | 0.3250 | 0.3170 | 0.5930 | 0.5680 | 0.4500 |
| SA3G | 0.4890 | 0.4890 | 0.4890 | 0.5150 | 0.5450 | 0.5110 | 0.5110 | 0.5110 | 0.5400 | 0.5400 | 0.5230 | 0.4850 | 0.4850 | 0.4670 | 0.4770 | 0.4780 | 0.4220 | 0.3770 | 0.4200 | 0.3830 |
| TP | 0.4690 | 0.4450 | 0.4450 | 0.5940 | 0.5780 | 0.5350 | 0.9050 | 0.9050 | 0.5450 | 0.5450 | 0.5240 | 0.4490 | 0.4490 | 0.4410 | 0.4170 | 0.4200 | 0.3710 | 0.4390 | 0.4210 | 0.4170 |
| DL4215 | 0.4960 | 0.4490 | 0.4490 | 0.5040 | 0.4740 | 0.5080 | 0.4440 | 0.4440 | 0.5440 | 0.5440 | 0.4810 | 0.4560 | 0.4560 | 0.4640 | 0.4550 | 0.4110 | 0.4230 | 0.3480 | 0.3480 | 0.3590 |
| 1587 | 0.5070 | 0.4810 | 0.4810 | 0.4540 | 0.4310 | 0.4230 | 0.3970 | 0.3970 | 0.4630 | 0.4630 | 0.5380 | 0.4810 | 0.4810 | 0.5260 | 0.5220 | 0.5960 | 0.6200 | 0.3660 | 0.3660 | 0.3440 |
| SL6Y | 0.4620 | 0.4440 | 0.4440 | 1.0000 | 0.7990 | 0.7370 | 0.5940 | 0.5940 | 0.4960 | 0.4960 | 0.4620 | 0.4200 | 0.4200 | 0.4360 | 0.4110 | 0.4200 | 0.4050 | 0.4340 | 0.4050 | 0.4480 |
| DRC186 | 0.4710 | 0.5070 | 0.5070 | 0.4140 | 0.4360 | 0.4320 | 0.4180 | 0.4180 | 0.4130 | 0.4130 | 0.4240 | 0.5110 | 0.5110 | 0.5220 | 0.5570 | 0.4810 | 0.4560 | 0.3620 | 0.3750 | 0.4130 |
| SA5Y | 0.4810 | 0.4890 | 0.4890 | 0.5080 | 0.5000 | 0.5190 | 0.5220 | 0.5220 | 0.5660 | 0.5660 | 0.5460 | 0.4930 | 0.4930 | 0.5040 | 0.4700 | 0.4400 | 0.4300 | 0.3720 | 0.3920 | 0.3750 |
| V52 | 0.4970 | 0.4810 | 0.4810 | 0.4690 | 0.4730 | 0.5000 | 0.5240 | 0.5240 | 0.5180 | 0.5180 | 0.9940 | 0.4110 | 0.4050 | 0.4850 | 0.4810 | 0.4920 | 0.4850 | 0.3770 | 0.3640 | 0.3470 |
| W6G | 1.0000 | 0.5830 | 0.5830 | 0.4620 | 0.4550 | 0.4500 | 0.4450 | 0.4450 | 0.4170 | 0.4170 | 0.4900 | 0.4830 | 0.4770 | 0.5360 | 0.5520 | 0.4530 | 0.4450 | 0.3810 | 0.3510 | 0.3700 |
| SA7G | 0.5830 | 1.0000 | 1.0000 | 0.4440 | 0.4810 | 0.4700 | 0.4220 | 0.4220 | 0.4930 | 0.4930 | 0.4740 | 0.5580 | 0.5580 | 0.5430 | 0.5510 | 0.5180 | 0.4890 | 0.3700 | 0.3410 | 0.3880 |
| SO5Y | 1.0000 | 0.5830 | 0.5830 | 0.4620 | 0.4550 | 0.4500 | 0.4450 | 0.4450 | 0.4170 | 0.4170 | 0.5000 | 0.4830 | 0.4770 | 0.5360 | 0.5520 | 0.4530 | 0.4450 | 0.3810 | 0.3510 | 0.3700 |
| SIO | 0.7330 | 0.6080 | 0.6080 | 0.4840 | 0.4840 | 0.4800 | 0.4390 | 0.4390 | 0.4410 | 0.4410 | 0.4720 | 0.5280 | 0.5280 | 0.5690 | 0.5600 | 0.4880 | 0.4800 | 0.3920 | 0.3610 | 0.3870 |
| W10G | 0.5410 | 0.5480 | 0.5480 | 0.4610 | 0.4770 | 0.4920 | 0.4370 | 0.4370 | 0.4890 | 0.4890 | 0.5380 | 0.5470 | 0.5470 | 0.5750 | 0.8140 | 0.4960 | 0.5110 | 0.3740 | 0.3820 | 0.3650 |
| SL5Y | 0.4620 | 0.4440 | 0.4440 | 1.0000 | 0.7990 | 0.7370 | 0.5940 | 0.5940 | 0.4960 | 0.4960 | 0.4620 | 0.4200 | 0.4200 | 0.4360 | 0.4110 | 0.4200 | 0.4050 | 0.4340 | 0.4050 | 0.4480 |
| SL4G | 0.4810 | 0.4890 | 0.4890 | 0.5080 | 0.5000 | 0.5190 | 0.5220 | 0.5220 | 0.5660 | 0.5660 | 0.5460 | 0.4930 | 0.4930 | 0.5040 | 0.4700 | 0.4400 | 0.4300 | 0.3720 | 0.3920 | 0.3750 |
| W7G | 1.0000 | 0.5830 | 0.5830 | 0.4620 | 0.4550 | 0.4500 | 0.4450 | 0.4450 | 0.4170 | 0.4170 | 0.4900 | 0.4830 | 0.4770 | 0.5360 | 0.5520 | 0.4530 | 0.4450 | 0.3810 | 0.3510 | 0.3700 |
| SA10G | 0.5830 | 1.0000 | 1.0000 | 0.4440 | 0.4810 | 0.4700 | 0.4220 | 0.4220 | 0.4930 | 0.4930 | 0.4740 | 0.5580 | 0.5580 | 0.5430 | 0.5510 | 0.5180 | 0.4890 | 0.3700 | 0.3410 | 0.3880 |
| E7G | 0.5830 | 1.0000 | 1.0000 | 0.4440 | 0.4810 | 0.4700 | 0.4220 | 0.4220 | 0.4930 | 0.4930 | 0.4740 | 0.5580 | 0.5580 | 0.5430 | 0.5510 | 0.5180 | 0.4890 | 0.3700 | 0.3410 | 0.3880 |
| DL4211 | 0.4620 | 0.4440 | 0.4440 | 1.0000 | 0.7990 | 0.7370 | 0.5940 | 0.5940 | 0.4960 | 0.4960 | 0.4620 | 0.4200 | 0.4200 | 0.4360 | 0.4110 | 0.4200 | 0.4050 | 0.4340 | 0.4050 | 0.4480 |
| VCSR05 | 0.4550 | 0.4810 | 0.4810 | 0.7990 | 1.0000 | 0.7740 | 0.5700 | 0.5700 | 0.5190 | 0.5190 | 0.4650 | 0.4730 | 0.4730 | 0.4810 | 0.4500 | 0.4270 | 0.4050 | 0.4340 | 0.3970 | 0.4480 |
| VCSR0207 | 0.4500 | 0.4700 | 0.4700 | 0.7370 | 0.7740 | 1.0000 | 0.5510 | 0.5510 | 0.5390 | 0.5390 | 0.5080 | 0.4540 | 0.4540 | 0.4320 | 0.4260 | 0.4150 | 0.4150 | 0.4530 | 0.3920 | 0.4110 |
| VCSR096 | 0.4450 | 0.4220 | 0.4220 | 0.5940 | 0.5700 | 0.5510 | 1.0000 | 1.0000 | 0.5370 | 0.5370 | 0.5160 | 0.4570 | 0.4570 | 0.4490 | 0.4250 | 0.4350 | 0.3940 | 0.4550 | 0.4300 | 0.4170 |
| V130003 | 0.4450 | 0.4220 | 0.4220 | 0.5940 | 0.5700 | 0.5510 | 1.0000 | 1.0000 | 0.5370 | 0.5370 | 0.5160 | 0.4570 | 0.4570 | 0.4490 | 0.4250 | 0.4350 | 0.3940 | 0.4550 | 0.4300 | 0.4170 |
| Env390 | 0.4170 | 0.4930 | 0.4930 | 0.4960 | 0.5190 | 0.5390 | 0.5370 | 0.5370 | 1.0000 | 1.0000 | 0.5260 | 0.4350 | 0.4350 | 0.4670 | 0.4550 | 0.4560 | 0.4530 | 0.3920 | 0.3980 | 0.3740 |
| Env9 | 0.4170 | 0.4930 | 0.4930 | 0.4960 | 0.5190 | 0.5390 | 0.5370 | 0.5370 | 1.0000 | 1.0000 | 0.5260 | 0.4350 | 0.4350 | 0.4670 | 0.4550 | 0.4560 | 0.4530 | 0.3920 | 0.3980 | 0.3740 |
| VCSR063 | 0.4900 | 0.4740 | 0.4740 | 0.4620 | 0.4650 | 0.5080 | 0.5160 | 0.5160 | 0.5260 | 0.5260 | 1.0000 | 0.4040 | 0.3990 | 0.4780 | 0.4730 | 0.4850 | 0.4920 | 0.3690 | 0.3570 | 0.3390 |
| VCSR051 | 0.4830 | 0.5580 | 0.5580 | 0.4200 | 0.4730 | 0.4540 | 0.4570 | 0.4570 | 0.4350 | 0.4350 | 0.4040 | 1.0000 | 0.9940 | 0.6070 | 0.5940 | 0.4810 | 0.4520 | 0.3210 | 0.3360 | 0.3590 |
| VCSR0102 | 0.4770 | 0.5580 | 0.5580 | 0.4200 | 0.4730 | 0.4540 | 0.4570 | 0.4570 | 0.4350 | 0.4350 | 0.3990 | 0.9940 | 1.0000 | 0.6070 | 0.5940 | 0.4810 | 0.4520 | 0.3210 | 0.3360 | 0.3590 |
| VCSR0162 | 0.5360 | 0.5430 | 0.5430 | 0.4360 | 0.4810 | 0.4320 | 0.4490 | 0.4490 | 0.4670 | 0.4670 | 0.4780 | 0.6070 | 0.6070 | 1.0000 | 0.6000 | 0.5040 | 0.4890 | 0.3460 | 0.3810 | 0.3690 |
| RFB16 | 0.5520 | 0.5510 | 0.5510 | 0.4110 | 0.4500 | 0.4260 | 0.4250 | 0.4250 | 0.4550 | 0.4550 | 0.4730 | 0.5940 | 0.5940 | 0.6000 | 1.0000 | 0.5150 | 0.5300 | 0.3710 | 0.3790 | 0.3780 |
| VCSR017 | 0.4530 | 0.5180 | 0.5180 | 0.4200 | 0.4270 | 0.4150 | 0.4350 | 0.4350 | 0.4560 | 0.4560 | 0.4850 | 0.4810 | 0.4810 | 0.5040 | 0.5150 | 1.0000 | 0.7720 | 0.3410 | 0.3260 | 0.3410 |
| NCTC30 | 0.4450 | 0.4890 | 0.4890 | 0.4050 | 0.4050 | 0.4150 | 0.3940 | 0.3940 | 0.4530 | 0.4530 | 0.4920 | 0.4520 | 0.4520 | 0.4890 | 0.5300 | 0.7720 | 1.0000 | 0.3480 | 0.3180 | 0.3250 |
| VCSR077 | 0.3810 | 0.3700 | 0.3700 | 0.4340 | 0.4340 | 0.4530 | 0.4550 | 0.4550 | 0.3920 | 0.3920 | 0.3690 | 0.3210 | 0.3210 | 0.3460 | 0.3710 | 0.3410 | 0.3480 | 1.0000 | 0.6010 | 0.4580 |
| strain20000 | 0.3510 | 0.3410 | 0.3410 | 0.4050 | 0.3970 | 0.3920 | 0.4300 | 0.4300 | 0.3980 | 0.3980 | 0.3570 | 0.3360 | 0.3360 | 0.3810 | 0.3790 | 0.3260 | 0.3180 | 0.6010 | 1.0000 | 0.4800 |
| VCSR045 | 0.3700 | 0.3880 | 0.3880 | 0.4480 | 0.4480 | 0.4110 | 0.4170 | 0.4170 | 0.3740 | 0.3740 | 0.3390 | 0.3590 | 0.3590 | 0.3690 | 0.3780 | 0.3410 | 0.3250 | 0.4580 | 0.4800 | 1.0000 |

**Fig. S9. Comparison of PilA proteins.** The heat map displaying fraction identity (fraction of identical residues) of PilA from patient and environmental isolates used in this study, which includes A1552 as a pandemic representative and *V. mimicus* (ATCC33655) as the outgroup. Green indicates fraction identity of 1, yellow indicates fraction identity >0.75, red fraction identity >0.5, and dark red fraction identity ≤0.5. The experimentally proven ability of the PilA variants to self-interact is indicated above the heatmap.

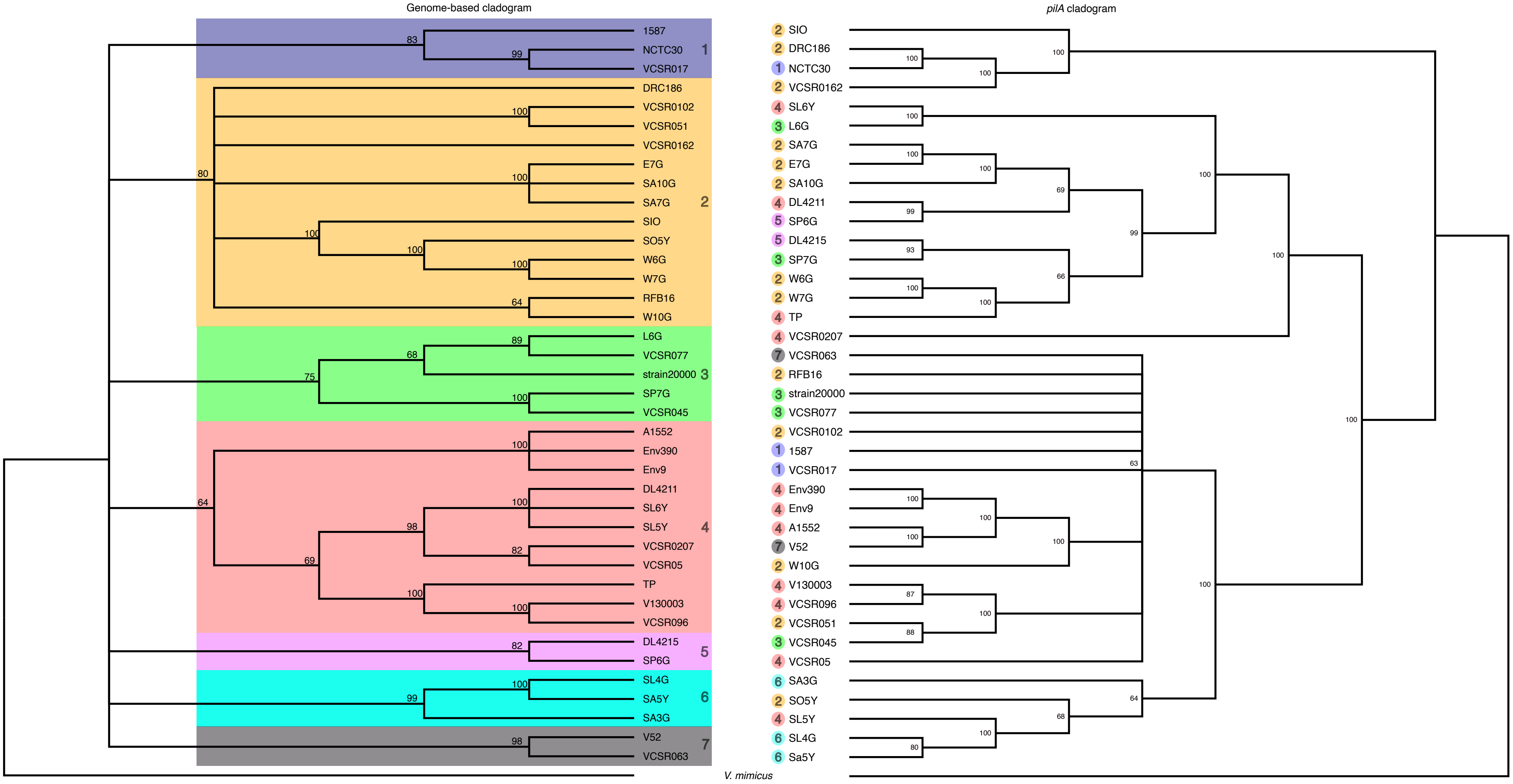


**Fig. S10. The *pilA* gene likely moves by horizontal gene transfer.** The figure compared the core gene-based cladogram of the 39 *V. cholerae* strains studied (on the left side) with the *pilA* nucleotide sequences-based cladogram (on the right side). Coloured boxes highlight supported clades of relatively related strains, which are also represented by coloured circles in the *pilA* cladogram. The incongruence between the genome-based and *pilA* reconstructions suggests horizontal gene transfer of *pilA*. *V. mimicus* (ATCC33655) is used as an outgroup. Statistical significance was verified using 100 bootstraps, and nodes with bootstrap values below 60 are collapsed.

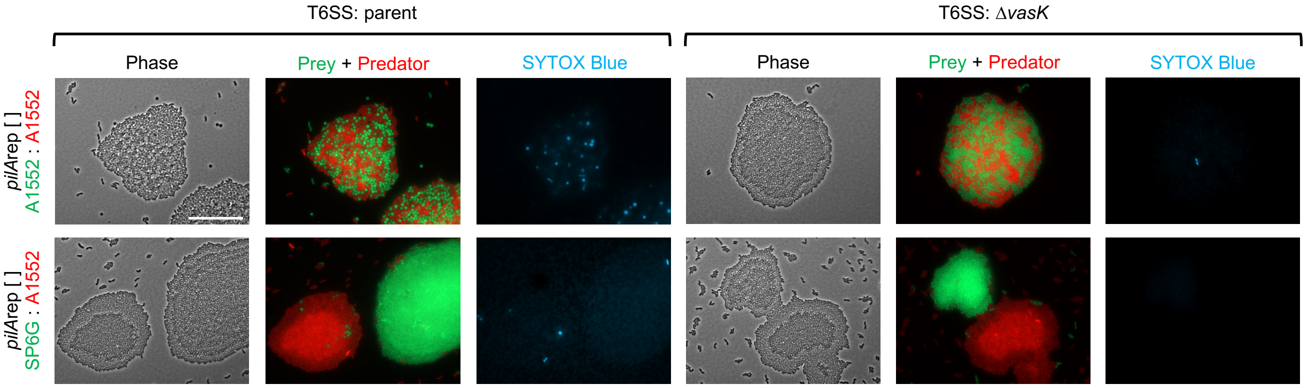


**Fig. S11. Pilus diversity provides T6SS protection by spatial segregation.** Microscopy images of co-culture experiments between T6SS-competent (Parent) or non-functional (Δ*vasK*) predator strains, and T6SS-sensitive (Δ4E/I) prey strains. Predator and prey strains are either carrying matching (upper panels), or diverse (lower panels) PilA variants. Phase contrast, a merge of prey (sfGFP, in green) and predator strains (mCherry, in red), and SYTOX Blue dead cell stain (blue) channels are displayed. Scalebar indicates 25 µm.

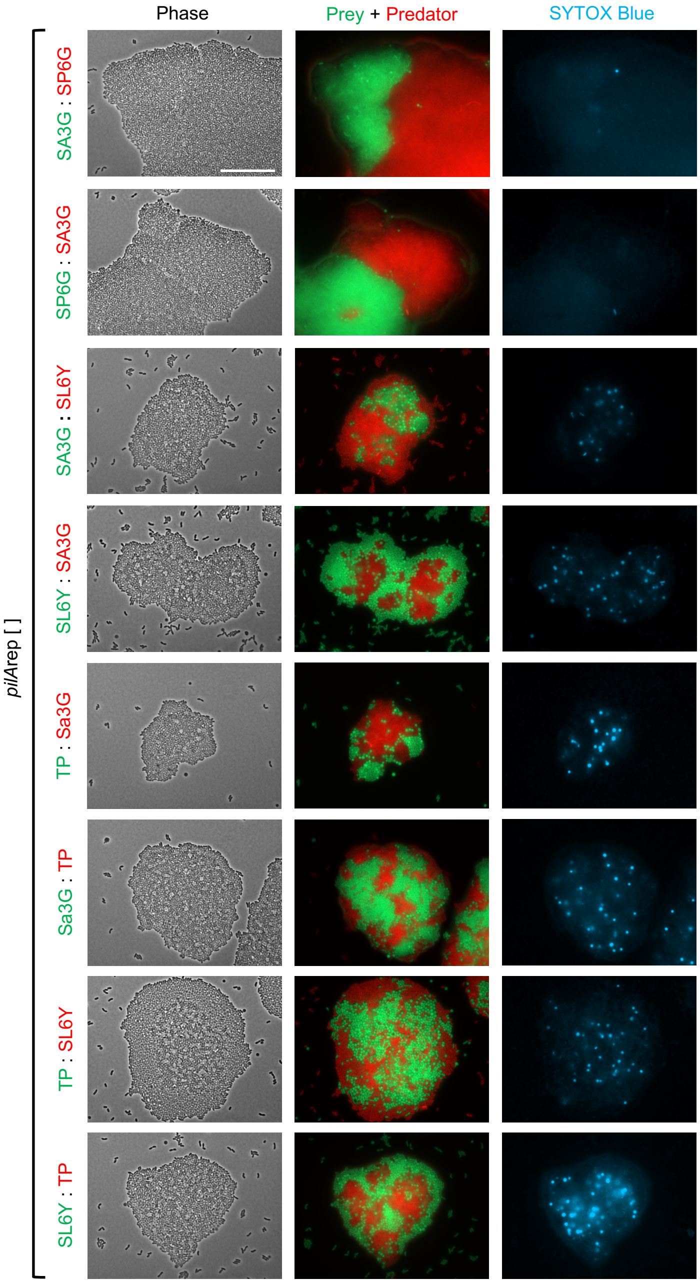


**Fig. S12. Microscopy of strain pairs with promiscuous T4P combinations.** Images of co-culture experiments between T6SS-competent (predator) and T6SS-sensitive (prey) strains, displaying all PilA variants, known to exhibit levels of promiscuousness. Phase contrast, a merge of prey (sfGFP, green) and predator strains (mCherry, red), and SYTOX Blue dead cell stain (blue) channels are displayed. Scalebar indicates 25 µm.

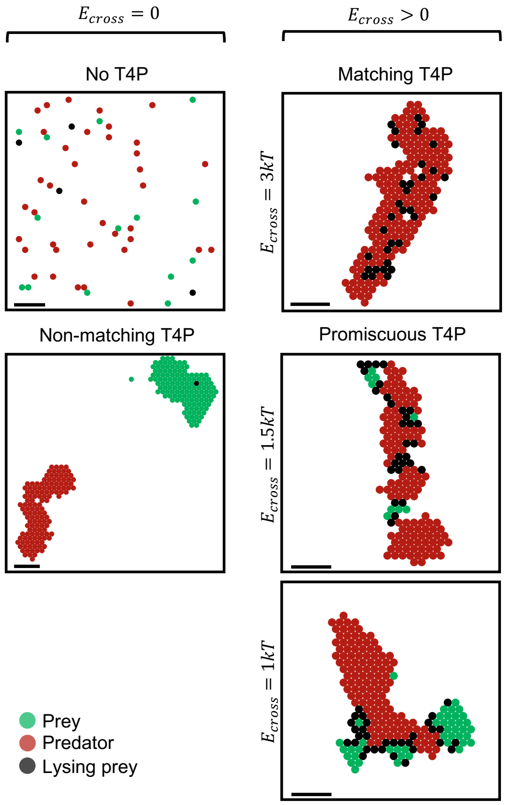


**Fig. S13. 2D simulations yield a similar phenomenology as 3D simulations.** In addition to the 3D simulations (see methods section), 2D simulations were performed in a triangular lattice, with the same division rate, diffusion coefficient, binding energies, total firing rate, and lysis rate as in the 3D case. The 5 panels show zooms from snapshots taken after 1 h of 2D simulations where 100x100 triangular lattices were initialized with 50 prey (green markers) and 50 predators (red markers) placed uniformly at random (note that this yields a larger density than in our 3D simulations). Lysing cells are represented by black markers. The black scale bar indicates the length of 5 marker diameters. On the left: no interaction between prey and predators ($E_{cross} = 0$). The top panel illustrates the complete absence of T4P, while the bottom panel corresponds to the non-matching T4P case. On the right: prey and predators interact via T4P with a non-zero binding energy ($E_{cross} > 0$). The top panel shows the matching T4P case, and the middle and bottom panels correspond to promiscuous T4P, with $E_{cross}$ equal to $1.5kT$ and $1kT$, respectively.

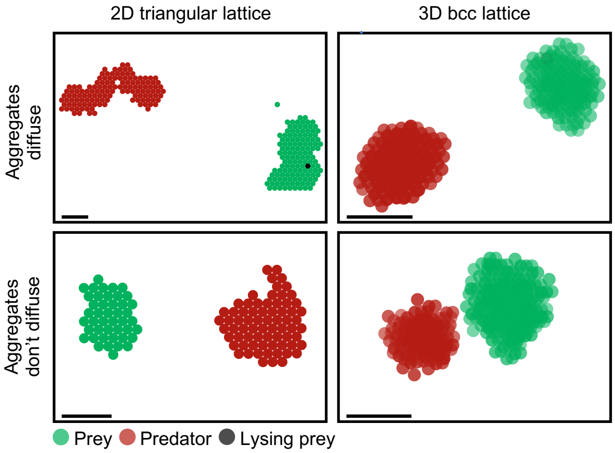


**Fig. S14. A similar phenomenology is obtained by four variants of the model.** On the left are 2D simulations, and on the right are 3D simulations. The top panels illustrate simulations that include the diffusion of aggregates as a whole, while the bottom panels depict simulations conducted without such diffusion (see methods). All panels show zooms from snapshots taken after 1 h of simulations performed as described in the methods section for 3D simulations, and as described in the legend of Figure S13 for 2D simulations. Prey and predators are represented by green and red markers, respectively, while lysing cells are represented by black markers. The black scale bar indicates the length of 5 marker diameters. The main behaviours (aggregation or no aggregation) remain consistent across all variants of the model. Notably, when aggregates do not diffuse, multiple prey and predator aggregates are formed after one hour. In contrast, when they diffuse, aggregates of a particular type coalesce, resulting in a single aggregate of prey and a single aggregate of predators.

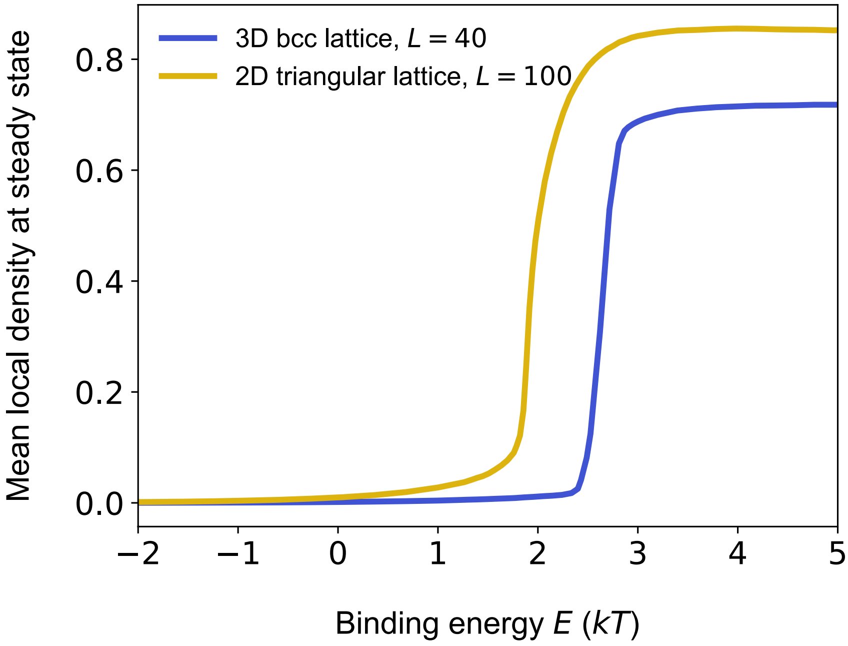


**Fig. S15. Aggregation is facilitated by T4P with sufficient binding energy.** The plot displays the mean local density, which represents the average fraction of occupied neighbouring sites of a bacterium, plotted against the T4P binding energy $E$ at steady state. In this figure, we consider a single type of bacteria (either prey or predators) diffusing and interacting through T4P with a specific attractive energy $E$, but without division or killing. The system can then be mapped to a lattice fluid model, and a Monte Carlo simulation of its steady state was performed. The blue curve represents the 3D model where bacteria diffuse on a $40\times40\times40$ body-centred cubic lattice, and the yellow curve corresponds to the 2D model where bacteria diffuse on a $100\times100$ triangular lattice. In both cases, the system comprises 100 bacteria.
