## Supplementary Table S1 for "Interactions between pili affect the outcome of bacterial competition driven by the type VI secretion system"

**Supplementary Table 1** – Bacterial strains and plasmids used in this study

| **Strain name** | **Strain #** | **Description** ^a, b^ | **Source** |
| --- | --- | --- | --- |
| ***Vibrio cholerae*** | | | |
| A1552 (WT) | GC#1 | A1552 Wild type, O1 El Tor Inaba; Rif^R^ | [1], genome sequence: [2] |
| A1552Δ*lacZ*::FRT | GC#1208 | A1552 deleted for *lacZ*; Rif^R^ | [3] |
| A1552-KanR-mCherry | GC#9700 | ∆VC1807::FRT-Kan^R^-FRT-PA1/04/03-mCherry_op13_; transcriptional fusion of Kan^R^ (*aph*)- and *mCherry*, which has been codon changed to enrich the A/T content in the first 13 codons of the 5’ region, template: [4]; *mCherry* is preceded by the constitutive *P*_lac_-derivative promoter, PA1/04/03; Rif^R^ | This study |
| A1552-SpecR-sfGFP | GC#9695 | ∆VC1807::FRT-Spec^R^-FRT-PA1/04/03-sfGFP; transcriptional fusion of Spec^R^ (*aad9*)- and *sfGFP*; *sfGFP* is preceded by the constitutive *P*_lac_-derivative promoter, PA1/04/03; Rif^R^ | This study |
| A1552-Tn*tfoX*, ∆*pilT* | GC#10495 | A1552-Tn*tfoX*, ∆*pilT*; Rif^R^ | This study |
| A1552-Tn*tfoX*, ∆*pilT*, ∆*pilA* | GC#10496 | A1552-Tn*tfoX*, ∆*pilT*, ∆*pilA*; Rif^R^ | This study |
| A1552-Tn*tfoX*, ∆*pilT*, ∆4E/I, SpecR-sfGFP | GC#10501 | Tn*tfoX*, ∆*pilT*, ΔVC1418-VC1419::FRT, ΔVCA0020-21::FRT, ΔVCA0123-24::FRT, ∆VCA0285-86::FRT, ∆VC1807::FRT-Spec^R^-FRT-PA1/04/03-sfGFP; Rif^R^ | This study |
| A1552-Tn*tfoX*, ∆*pilT*, KanR-mCherry | GC#10497 | Tn*tfoX*, ∆*pilT*, ∆VC1807::FRT-Kan^R^-FRT-PA1/04/03- mCherry_op13_; Rif^R^ | This study |
| A1552-Tn*tfoX*, ∆*pilT*, ∆*pilA*, ∆4E/I, SpecR-sfGFP | GC#10498 | Tn*tfoX*, ∆*pilT*, ∆*pilA*, ΔVC1418-VC1419::FRT, ΔVCA0020-21::FRT, ΔVCA0123-24::FRT, ∆VCA0285-86::FRT, ∆VC1807::FRT-Spec^R^-FRT-PA1/04/03-sfGFP; Rif^R^ | This study |
| A1552-Tn*tfoX*, ∆*pilT*, ∆*pilA*, KanR-mCherry | GC#10500 | Tn*tfoX*, ∆*pilT*, ∆*pilA*, ∆VC1807::FRT-Kan^R^-FRT-PA1/04/03-mCherry_op13_; RifR | This study |
| A1552-Tn*tfoX*, ∆*pilT*, ∆*vasK*, KanR-mCherry | GC#10499 | Tn*tfoX*, ∆*pilT*, ∆*vasK*, ∆VC1807::FRT-Kan^R^-FRT-PA1/04/03-mCherry_op13_; Rif^R^ | This study |
| A1552-Tn*tfoX*, ∆*pilT*, ∆*pilA*, ∆*vasK*, KanR-mCherry | GC#10507 | Tn*tfoX*, ∆*pilT*, ∆*pilA*, ∆*vasK*, ∆VC1807::FRT-Kan^R^-FRT-PA1/04/03-mCherry_op13_; Rif^R^ | This study |
| A1552-Tn*tfoX*, ∆*pilT*, *pilA*rep[A1552], ∆4E/I, SpecR-sfGFP | GC#10504 | Tn*tfoX*, ∆*pilT*, *pilA*rep[A1552], ΔVC1418-VC1419::FRT, ΔVCA0020-21::FRT, ΔVCA0123-24::FRT, ∆VCA0285-86::FRT, ∆VC1807::FRT-Spec^R^-FRT-PA1/04/03-sfGFP; Rif^R^ | This study |
| A1552-Tn*tfoX*, ∆*pilT*, *pilA*rep[A1552], KanR-mCherry | GC#10508 | Tn*tfoX*, ∆*pilT*, *pilA*rep[A1552], ∆VC1807::FRT-Kan^R^-FRT-PA1/04/03-mCherry_op13_; Rif^R^ | This study |
| A1552-Tn*tfoX*, ∆*pilT*, *pilA*rep[SP6G], ∆4E/I, SpecR-sfGFP | GC#10505 | Tn*tfoX*, ∆*pilT*, *pilA*rep[SP6G], ΔVC1418-VC1419::FRT, ΔVCA0020-21::FRT, ΔVCA0123-24::FRT, ∆VCA0285-86::FRT, ∆VC1807::FRT-Spec^R^-FRT-PA1/04/03-sfGFP; Rif^R^ | This study |
| A1552-Tn*tfoX*, ∆*pilT*, *pilA*rep[SP6G], KanR-mCherry | GC#10522 | Tn*tfoX*, ∆*pilT*, *pilA*rep[SP6G], ∆VC1807::FRT-Kan^R^-FRT-PA1/04/03-mCherry_op13_; Rif^R^ | This study |
| A1552-Tn*tfoX*, ∆*pilT*, *pilA*rep[SP7G], ∆4E/I, SpecR-sfGFP | GC#10510 | Tn*tfoX*, ∆*pilT*, *pilA*rep[SP7G], ΔVC1418-VC1419::FRT, ΔVCA0020-21::FRT, ΔVCA0123-24::FRT, ∆VCA0285-86::FRT, ∆VC1807::FRT-Spec^R^-FRT-PA1/04/03-sfGFP; Rif^R^ | This study |
| A1552-Tn*tfoX*, ∆*pilT*, *pilA*rep[SP7G], KanR-mCherry | GC#10518 | Tn*tfoX*, ∆*pilT*, *pilA*rep[SP7G], ∆VC1807::FRT-Kan^R^-FRT-PA1/04/03-mCherry_op13_; Rif^R^ | This study |
| A1552-Tn*tfoX*, ∆*pilT*, *pilA*rep[L6G], ∆4E/I, SpecR-sfGFP | GC#10512 | Tn*tfoX*, ∆*pilT*, *pilA*rep[L6G], ΔVC1418-VC1419::FRT, ΔVCA0020-21::FRT, ΔVCA0123-24::FRT, ∆VCA0285-86::FRT, ∆VC1807::FRT-Spec^R^-FRT-PA1/04/03-sfGFP; Rif^R^ | This study |
| A1552-Tn*tfoX*, ∆*pilT*, *pilA*rep[L6G], KanR-mCherry | GC#10520 | Tn*tfoX*, ∆*pilT*, *pilA*rep[L6G], ∆VC1807::FRT-Kan^R^-FRT-PA1/04/03-mCherry_op13_; Rif^R^ | This study |
| A1552-Tn*tfoX*, ∆*pilT*, *pilA*rep[SA3G], ∆4E/I, SpecR-sfGFP | GC#10513 | Tn*tfoX*, ∆*pilT*, *pilA*rep[SA3G], ΔVC1418-VC1419::FRT, ΔVCA0020-21::FRT, ΔVCA0123-24::FRT, ∆VCA0285-86::FRT, ∆VC1807::FRT-Spec^R^-FRT-PA1/04/03-sfGFP; Rif^R^ | This study |
| A1552-Tn*tfoX*, ∆*pilT*, *pilA*rep[SA3G], KanR-mCherry | GC#10521 | Tn*tfoX*, ∆*pilT*, *pilA*rep[SA3G], ∆VC1807::FRT-Kan^R^-FRT-PA1/04/03-mCherry_op13_; Rif^R^ | This study |
| A1552-Tn*tfoX*, ∆*pilT*, *pilA*rep[TP], ∆4E/I, SpecR-sfGFP | GC#10514 | Tn*tfoX*, ∆*pilT*, *pilA*rep[TP], ΔVC1418-VC1419::FRT, ΔVCA0020-21::FRT, ΔVCA0123-24::FRT, ∆VCA0285-86::FRT, ∆VC1807::FRT-Spec^R^-FRT-PA1/04/03-sfGFP; Rif^R^ | This study |
| A1552-Tn*tfoX*, ∆*pilT*, *pilA*rep[TP], KanR-mCherry | GC#10523 | Tn*tfoX*, ∆*pilT*, *pilA*rep[TP], ∆VC1807::FRT-Kan^R^-FRT-PA1/04/03-mCherry_op13_; Rif^R^ | This study |
| A1552-Tn*tfoX*, ∆*pilT*, *pilA*rep[DL4215], ∆4E/I, SpecR-sfGFP | GC#10515 | Tn*tfoX*, ∆*pilT*, *pilA*rep[DL4215], ΔVC1418-VC1419::FRT, ΔVCA0020-21::FRT, ΔVCA0123-24::FRT, ∆VCA0285-86::FRT, ∆VC1807::FRT-Spec^R^-FRT-PA1/04/03-sfGFP; Rif^R^ | This study |
| A1552-Tn*tfoX*, ∆*pilT*, *pilA*rep[DL4215], KanR-mCherry | GC#10524 | Tn*tfoX*, ∆*pilT*, *pilA*rep[DL4215], ∆VC1807::FRT-Kan^R^-FRT-PA1/04/03-mCherry_op13_; Rif^R^ | This study |
| A1552-Tn*tfoX*, ∆*pilT*, *pilA*rep[1587], ∆4E/I, SpecR-sfGFP | GC#10516 | Tn*tfoX*, ∆*pilT*, *pilA*rep[1587], ΔVC1418-VC1419::FRT, ΔVCA0020-21::FRT, ΔVCA0123-24::FRT, ∆VCA0285-86::FRT, ∆VC1807::FRT-Spec^R^-FRT-PA1/04/03-sfGFP; Rif^R^ | This study |
| A1552-Tn*tfoX*, ∆*pilT*, *pilA*rep[1587], KanR-mCherry | GC#10525 | Tn*tfoX*, ∆*pilT*, *pilA*rep[1587], ∆VC1807::FRT-Kan^R^-FRT-PA1/04/03-mCherry_op13_; Rif^R^ | This study |
| A1552-Tn*tfoX*, ∆*pilT*, *pilA*rep[SL6Y], ∆4E/I, SpecR-sfGFP | GC#10511 | Tn*tfoX*, ∆*pilT*, *pilA*rep[SL6Y], ΔVC1418-VC1419::FRT, ΔVCA0020-21::FRT, ΔVCA0123-24::FRT, ∆VCA0285-86::FRT, ∆VC1807::FRT-Spec^R^-FRT-PA1/04/03-sfGFP; Rif^R^ | This study |
| A1552-Tn*tfoX*, ∆*pilT*, *pilA*rep[SL6Y], KanR-mCherry | GC#10519 | Tn*tfoX*, ∆*pilT*, *pilA*rep[SL6Y], ∆VC1807::FRT-Kan^R^-FRT-PA1/04/03-mCherry_op13_; Rif^R^ | This study |
| A1552-Tn*tfoX*, ∆*pilT*, *pilA*rep[DRC186], ∆4E/I, SpecR-sfGFP | GC#10517 | Tn*tfoX*, ∆*pilT*, *pilA*rep[DRC186], ΔVC1418-VC1419::FRT, ΔVCA0020-21::FRT, ΔVCA0123-24::FRT, ∆VCA0285-86::FRT, ∆VC1807::FRT-Spec^R^-FRT-PA1/04/03-sfGFP; Rif^R^ | This study |
| A1552-Tn*tfoX*, ∆*pilT*, *pilA*rep[DRC186], KanR-mCherry | GC#10526 | Tn*tfoX*, ∆*pilT*, *pilA*rep[DRC186], ∆VC1807::FRT-Kan^R^-FRT-PA1/04/03-mCherry_op13_; Rif^R^ | This study |
| A1552-Tn*tfoX*, ∆*pilT*, *pilA*rep[SA5Y], ∆4E/I, SpecR-sfGFP | GC#10506 | Tn*tfoX*, ∆*pilT*, *pilA*rep[SA5Y], ΔVC1418-VC1419::FRT, ΔVCA0020-21::FRT, ΔVCA0123-24::FRT, ∆VCA0285-86::FRT, ∆VC1807::FRT-Spec^R^-FRT-PA1/04/03-sfGFP; Rif^R^ | This study |
| A1552-Tn*tfoX*, ∆*pilT*, *pilA*rep[SA5Y], KanR-mCherry | GC#10528 | Tn*tfoX*, ∆*pilT*, *pilA*rep[SA5Y], ∆VC1807::FRT-Kan^R^-FRT-PA1/04/03-mCherry_op13_; Rif^R^ | This study |
| A1552-Tn*tfoX*, ∆*pilT*, *pilA*rep[V52], ∆4E/I, SpecR-sfGFP | GC#10535 | Tn*tfoX*, ∆*pilT*, *pilA*rep[V52], ΔVC1418-VC1419::FRT, ΔVCA0020-21::FRT, ΔVCA0123-24::FRT, ∆VCA0285-86::FRT, ∆VC1807::FRT-Spec^R^-FRT-PA1/04/03-sfGFP; Rif^R^ | This study |
| A1552-Tn*tfoX*, ∆*pilT*, *pilA*rep[V52], KanR-mCherry | GC#10529 | Tn*tfoX*, ∆*pilT*, *pilA*rep[V52], ∆VC1807::FRT-Kan^R^-FRT-PA1/04/03-mCherry_op13_; Rif^R^ | This study |
| A1552-Tn*tfoX*, ∆*pilT*, *pilA*rep[W6G], ∆4E/I, SpecR-sfGFP | GC#10536 | Tn*tfoX*, ∆*pilT*, *pilA*rep[W6G], ΔVC1418-VC1419::FRT, ΔVCA0020-21::FRT, ΔVCA0123-24::FRT, ∆VCA0285-86::FRT, ∆VC1807::FRT-Spec^R^-FRT-PA1/04/03-sfGFP; Rif^R^ | This study |
| A1552-Tn*tfoX*, ∆*pilT*, *pilA*rep[W6G], KanR-mCherry | GC#10530 | Tn*tfoX*, ∆*pilT*, *pilA*rep[W6G], ∆VC1807::FRT-Kan^R^-FRT-PA1/04/03-mCherry_op13_; Rif^R^ | This study |
| A1552-Tn*tfoX*, ∆*pilT*, *pilA*rep[SA7G], ∆4E/I, SpecR-sfGFP | GC#10537 | Tn*tfoX*, ∆*pilT*, *pilA*rep[SA7G], ΔVC1418-VC1419::FRT, ΔVCA0020-21::FRT, ΔVCA0123-24::FRT, ∆VCA0285-86::FRT, ∆VC1807::FRT-Spec^R^-FRT-PA1/04/03-sfGFP; Rif^R^ | This study |
| A1552-Tn*tfoX*, ∆*pilT*, *pilA*rep[SA7G], KanR-mCherry | GC#10531 | Tn*tfoX*, ∆*pilT*, *pilA*rep[SA7G], ∆VC1807::FRT-Kan^R^-FRT-PA1/04/03-mCherry_op13_; Rif^R^ | This study |
| A1552-Tn*tfoX*, ∆*pilT*, *pilA*rep[SO5Y], ∆4E/I, SpecR-sfGFP | GC#10538 | Tn*tfoX*, ∆*pilT*, *pilA*rep[SO5Y], ΔVC1418-VC1419::FRT, ΔVCA0020-21::FRT, ΔVCA0123-24::FRT, ∆VCA0285-86::FRT, ∆VC1807::FRT-Spec^R^-FRT-PA1/04/03-sfGFP; Rif^R^ | This study |
| A1552-Tn*tfoX*, ∆*pilT*, *pilA*rep[SO5Y], KanR-mCherry | GC#10532 | Tn*tfoX*, ∆*pilT*, *pilA*rep[SO5Y], ∆VC1807::FRT-Kan^R^-FRT-PA1/04/03-mCherry_op13_; Rif^R^ | This study |
| A1552-Tn*tfoX*, ∆*pilT*, *pilA*rep[SIO], ∆4E/I, SpecR-sfGFP | GC#10527 | Tn*tfoX*, ∆*pilT*, *pilA*rep[SIO], ΔVC1418-VC1419::FRT, ΔVCA0020-21::FRT, ΔVCA0123-24::FRT, ∆VCA0285-86::FRT, ∆VC1807::FRT-Spec^R^-FRT-PA1/04/03-sfGFP; Rif^R^ | This study |
| A1552-Tn*tfoX*, ∆*pilT*, *pilA*rep[SIO], KanR-mCherry | GC#10533 | Tn*tfoX*, ∆*pilT*, *pilA*rep[SIO], ∆VC1807::FRT-Kan^R^-FRT-PA1/04/03-mCherry_op13_; Rif^R^ | This study |
| A1552-Tn*tfoX*, ∆*pilT*, *pilA*rep[W10G], ∆4E/I, SpecR-sfGFP | GC#10539 | Tn*tfoX*, ∆*pilT*, *pilA*rep[W10G], ΔVC1418-VC1419::FRT, ΔVCA0020-21::FRT, ΔVCA0123-24::FRT, ∆VCA0285-86::FRT, ∆VC1807::FRT-Spec^R^-FRT-PA1/04/03-sfGFP; Rif^R^ | This study |
| A1552-Tn*tfoX*, ∆*pilT*, *pilA*rep[W10G], KanR-mCherry | GC#10534 | Tn*tfoX*, ∆*pilT*, *pilA*rep[W10G], ∆VC1807::FRT-Kan^R^-FRT-PA1/04/03-mCherry_op13_; Rif^R^ | This study |
| A1552-Tn*tfoX*, ∆*pilT*, *pilA*rep[A1552], ∆*vasK*, KanR-mCherry | GC#10509 | Tn*tfoX*, ∆*pilT*, *pilA*rep[A1552], ∆*vasK*, ∆VC1807::FRT-Kan^R^-FRT-PA1/04/03-mCherry_op13_; Rif^R^ | This study |
| ***Escherichia coli*** |  |  |  |
| S17-1 λpir | GC#648 | TpR Sm^R^ *recA thi pro hsdR-M+ RP4:2-Tc:Mu:*Kan^R^ Tn*7* (λ*pir*) | [5] |
| SM10 λpir | GC#647 | *thi* *thr* *leu* *tonA* *lacY* *supE* *recA::RP4-2-Tc:Mu:*Kan^R^ (λ*pir*) | [5] |
| **Plasmids** | | | |
| pUX-BF13 | GC#457 | pUX-BF13 - oriR6K, helper plasmid with Tn7 transposition function; Amp^R^ | [6] |
| pGP704-mTn*tfoX* | GC#1624 | pGP704 with mini-Tn7 carrying *araC* and *P*_BAD_-*tfoX*; Amp^R^, Gent^R^ (Tn*tfoX*) | [7] |
| pGP704-Sac28 (p28) | GC#649 | pGP704-Sac28 suicide vector, *ori*R6K, *sacB*; Amp^R^ | [8] |
| p28-*pilA* | GC#1050 | pGP704-Sac28-∆*pilA* | [8] |
| p28-∆*pilT* | GC#5959 | pGP704-Sac28-∆*pilT* | [9] |
| p28-*pilA*rep[A1552] | GC#6199 | pGP704-Sac28-*pilA*rep[A1552] | [9] |
| p28-*pilA*rep[SP6G] | GC#5674 | pGP704-Sac28-*pilA*rep[964; SP6G] | [9] |
| p28-*pilA*rep[SP7G] | GC#5665 | pGP704-Sac28-*pilA*rep[952; SP7G] | [9] |
| p28-*pilA*rep[L6G] | GC#5666 | pGP704-Sac28-*pilA*rep[956; L6G] | [9] |
| p28-*pilA*rep[SA3G] | GC#5667 | pGP704-Sac28-*pilA*rep[957; SA3G] | [9] |
| p28-*pilA*rep[TP] | GC#5670 | pGP704-Sac28-*pilA*rep[999; TP] | [9] |
| p28-*pilA*rep[DL4215] | GC#5671 | pGP704-Sac28-*pilA*rep[3079; DL4215] | [9] |
| p28-*pilA*rep[1587] | GC#5672 | pGP704-Sac28-*pilA*rep[3081; 1587] | [9] |
| p28-*pilA*rep[SL6Y] | GC#5673 | pGP704-Sac28-*pilA*rep[953; SL6Y] | [9] |
| p28-*pilA*rep[DRC186] | GC#5676 | pGP704-Sac28-*pilA*rep[2501; DRC186] | [9] |
| p28-*pilA*rep[SA5Y] | GC#6200 | pGP704-Sac28-*pilA*rep[353; SA5Y] | [9] |
| p28-*pilA*rep[V52] | GC#6201 | pGP704-Sac28-*pilA*rep[1510; V52] | [9] |
| p28-*pilA*rep[W6G] | GC#5664 | pGP704-Sac28-*pilA*rep[354; W6G] | [9] |
| p28-*pilA*rep[SA7G] | GC#5668 | pGP704-Sac28-*pilA*rep[959; SA7G] | [9] |
| p28-*pilA*rep[SO5Y] | GC#5669 | pGP704-Sac28-*pilA*rep[960; SO5Y] | [9] |
| p28-*pilA*rep[SIO] | GC#5675 | pGP704-Sac28-*pilA*rep[1000; SIO] | [9] |
| p28-*pilA*rep[W10G] | GC#5677 | pGP704-Sac28-*pilA*rep[5537; W10G] | [9] |
| p28-∆VCA0285-86::FRT | GC#10540 | pGP704-Sac28-∆VCA0285-86::FRT | This study |
| p28-∆*vasK* | GC#1124 | pGP704-Sac28-ΔVCA0120 | [10] |
| p28-KanR-mCherry | GC#10542 | pGP704-Sac28-∆VC1807::FRT-Kan^R^-FRT-PA1/04/03-mCherry_op13_ | This study |
| p28-SpecR-sfGFP | GC#10541 | pGP704-Sac28-∆VC1807::FRT-Spec^R^-FRT-PA1/04/03-sfGFP | This study |

^a^ Strains containing mini-Tn7 insertions with arabinose-inducible constructs (Tn*tfoX*) are Gent^R^

^b^ FRT; flippase recognition target (FRT), if only one: scar left behind by TransFLP method [3,11,12]
