## Supplementary Table S2 for "Interactions between pili affect the outcome of bacterial competition driven by the type VI secretion system"

**Supplementary Table 2** – Information on genomes used for bioinformatical analysis.

| **Strain name** | **Strain #** | **Description** | **Source** | **Version** |
| --- | --- | --- | --- | --- |
| ***Vibrio mimicus*** | | | |  |
| ATCC33655 | N.A. | Human isolate, TN, USA | unpublished | GCF_001471395.1 |
| ***Vibrio spp.***^a^ | | | |  |
| A1552 | GC#1 | Patient isolate, O1 serogroup, El Tor biotype, Inaba serotype, Peru | [1], genome sequence: [2] | GCA_003097695.1 |
| SP6G | GC#964 | Environmental isolate; non-O1/O139 serogroup, CA, USA | [3], genome sequence: [4] | GCA_013357745.1 |
| SP7G | GC#952 | Environmental isolate; non-O1/O139 serogroup, CA, USA | [3], genome sequence: [4] | GCA_013357765.1 |
| L6G | GC#956 | Environmental isolate; non-O1/O139 serogroup, CA, USA | [3], genome sequence: [4] | GCA_013357685.1 |
| SA3G | GC#957 | Environmental isolate; non-O1/O139 serogroup, CA, USA | [3], genome sequence: [4] | GCA_013357885.1 |
| TP | GC#999 | Environmental isolate; non-O1/O139 serogroup, CA, USA | [5] | NCBI-SRA, Accession number: SRX22109611 |
| DL4215 | GC#3079 | Environmental isolate; O113 serogroup, Rio Grande, USA | [6] | NCBI-SRA, Accession number: SRX22109613 |
| 1587 | GC#3081 | Patient isolate; O12 serogroup, Peru | [6] | NCBI-SRA, Accession number: SRX22109614 |
| SL6Y | GC#953 | Environmental isolate; non-O1/O139 serogroup, CA, USA | [3], genome sequence: [4] | GCA_013357725.1 |
| DRC186 | GC#2501 | Unknown isolate; non-O1 serogroup, Democratic Republic of the Congo | unpublished | GCA_030710345.1 |
| SA5Y | GC#353 | Environmental isolate; non-O1/O139 serogroup, CA, USA | [3], genome sequence: [2] | GCA_003063885.1 |
| V52 | GC#1510 | Patient isolate; O37 serogroup, Sudan | [7,8] | NCBI-SRA, Accession number: SRX22139550 |
| W6G | GC#354 | Environmental isolate; non-O1/O139 serogroup, CA, USA | [3], genome sequence: [4] | GCA_013357785.1 |
| SA7G | GC#959 | Environmental isolate; non-O1/O139 serogroup, CA, USA | [3], genome sequence: [4] | GCA_013357845.1 |
| SO5Y | GC#960 | Environmental isolate; non-O1/O139 serogroup, CA, USA | [3], genome sequence: [4] | GCA_013357665.1 |
| SIO | GC#1000 | Environmental isolate; non-O1/O139 serogroup, CA, USA | [5] | NCBI-SRA, Accession number: SRX22109610 |
| W10G | GC#5537 | Environmental isolate; non-O1/O139 serogroup, CA, USA | [3], genome sequence: [4] | GCA_013357605.1 |
| SL5Y | GC#954 | Environmental isolate; non-O1/O139 serogroup, CA, USA | [3], genome sequence: [4] | GCA_013357645.1 |
| SL4G | GC#955 | Environmental isolate; non-O1/O139 serogroup, CA, USA | [3], genome sequence: [4] | GCA_013357625.1 |
| W7G | GC#962 | Environmental isolate; non-O1/O139 serogroup, CA, USA | [3], genome sequence: [4] | GCA_013357805.1 |
| SA10G | GC#5539 | Environmental isolate; non-O1/O139 serogroup, CA, USA | [3], genome sequence: [4] | GCA_013357865.1 |
| E7G | GC#963 | Environmental isolate; non-O1/O139 serogroup, CA, USA | [3], genome sequence: [4] | GCA_013357825.1 |
| DL4211 | GC#3084 | Environmental isolate; O123 serogroup, Rio Grande, USA | [6] | NCBI-SRA, Accession number: SRX22109612 |
| VCSR05 | N.A. | Patient isolate (diarrhea); O5 serogroup, Philippines | [9] | GCA_023168205.1 |
| VCSR0207 | N.A. | Environmental isolate; O207 serogroup, Japan | [9] | GCA_023168105.1 |
| VCSR096 | N.A. | Patient isolate (diarrhea); O96 serogroup, India | [9] | GCA_023168285.1 |
| V130003 | N.A. | Patient isolate (feces); O144 serogroup, Japan | unpublished | GCA_023169825.1 |
| Env390 | N.A. | Environmental isolate; non-toxigenic O1 serogroup, hybrid (El Tor/Classical) biotype, Ogawa serotype, Gressier, Haiti | [10], genome sequence: [11] | GCF_001854425.1 |
| Env9 | N.A. | Environmental isolate; non-toxigenic O1 serogroup, hybrid (El Tor/Classical) biotype, Ogawa serotype, La Salle, Haiti | [10], genome sequence: [11] | GCF_000788715.2 |
| VCSR063 | N.A. | Patient isolate (diarrhea); O63 serogroup, India | [9] | GCA_023168245.1 |
| VCSR051 | N.A. | Patient isolate (diarrhea); O51 serogroup, India | [9] | GCA_023168225.1 |
| VCSR0102 | N.A. | Patient isolate (diarrhea); O102 serogroup, China | [9] | GCA_023167925.1 |
| VCSR0162 | N.A. | Patient isolate (diarrhea); O162 serogroup, Argentina | [9] | GCA_023167985.1 |
| RFB16 | N.A. | Environmental isolate, unknown serogroup, North Park Lake (PA, USA) | [12] | GCA_008369605.1 |
| VCSR017 | N.A. | Patient isolate (diarrhea); O17 serogroup, India | [9] | GCA_023168045.1 |
| NCTC30 | N.A. | Patient isolate (diarrhea), probably O2 serogroup, Egypt | [13] | GCA_900538065.1 |
| VCSR077 | N.A. | Patient isolate (diarrhea); O77 serogroup, India | [9] | GCA_023168265.1 |
| strain20000 | N.A. | Environmental isolate; O1 serogroup, Rostov-on-Don, Russia | unpublished | GCA_004328575.1 |
| VCSR045 | N.A. | Patient isolate (diarrhea); O45 serogroup, India | [9] | GCA_023168185.1 |
| 2012EL-1759 | N.A. | Environmental; O395 serogroup, Haiti | [14], genome sequence: [15] | JNEW00000000.1 |
| HE45 | N.A. | Environmental isolate; unknown serogroup, Haiti | [6] | GCF_000279285.1 |
| A325 | N.A. | Unknown isolate; non-O1/O139 serogroup, Argentina | [14] | CWSO00000000.1 |
| 490−93 | N.A. | Patient isolate; O155 serogroup, Thailand | [14], genome sequence: [16] | JIDQ00000000.1 |
| TMA21 | N.A. | Environmental isolate; non-O1/O139 serogroup, Brazil | [6] | ACHY00000000 |
| MZO-2 | N.A. | Patient isolate; O14 serogroup, Bangladesh | [6] | GCA_000153985.3 |
| TM11079−80 | N.A. | Sewage isolate; non-toxigenic O1 El Tor serogroup, Brazil | [14] | GCA_000174255.1 |
| PG41 | N.A. | *Vibrio fluvialis*, Patient isolate; Bahrain | [14], genome sequence: [17] | ASXS00000000.1 |
| VL426 | N.A. | Environmental isolate; non-O1/O139 serogroup, Germany | [6] | GCF_000174235.1 |
| MZO-3 | N.A. | Patient isolate; O37 serogroup, Bangladesh | [6] | GCA_000168935.3 |
| AM-19226 | N.A. | Patient isolate; non-O1/O139 serogroup, Bangladesh | [18] | GCA_000153785.3 |
| S12 | N.A. | Environmental isolate; non-O1/O139 serogroup, Australia | [19] | MDST00000000 |
| BC1071 | N.A. | Patient isolate; unknown serogroup, Germany | [20] | LT897798.1 |

^a^ Unless stated otherwise, all *Vibrio spp.* strains are *Vibrio cholerae*
